# A mechanism for telomere-specific telomere length regulation

**DOI:** 10.1101/2024.06.12.598646

**Authors:** Gabriela M. Teplitz, Emeline Pasquier, Erin Bonnell, Evelina De Laurentiis, Louise Bartle, Jean-François Lucier, Dean S. Dawson, Raymund J. Wellinger

## Abstract

Telomere length is a critical determinant of telomere function and hence chromosome stability. Critically short telomeres induce cellular senescence and division arrest, which eventually may lead to devastating age-related degenerative diseases. Conversely, maintenance of telomere length is a hallmark of cancer. How telomere set-length is established and molecular mechanisms for telomere-specific length regulation remained unknown. Here we detail a mechanism of a telomere-specific set-length regulation that causes drastic differences in telomere length between individual telomeres in the same cell. Indeed, the results show that telomerase recruitment is modulated in *cis* in a telomere-specific way. Increased Sir4 abundance on yeast TEL03L subtelomeric heterochromatin leads to a set-length maintenance that is two to three times higher than on any other telomere. Remarkably, the distal 15 kb of TEL03L are sufficient to transfer this telomere specific set-length regulation to another chromosome. Furthermore, a mutation in the telomere boundary element protein Tbf1 allows increased Sir4 binding on all telomeres and hence results in longer set-lengths. The results therefore will force a rethinking of telomere length regulation away from the generalized view that all telomeres are treated the same to a more telomere-specific treatment.

**HIGHLIGHTS:**

- Regulation of the set-length of telomeric repeats is telomere-specific.
- TEL03L on yeast chromosome III displays a set-length regulation that confers an extremely long repeat tract.
- Transferring the distal part of TEL03L onto chromosome XV is sufficient to impose the very long set-length regulation.
- Telomere-specific tract set-length regulation depends on the alternate telomerase recruitment pathway involving Sir4 and yKU and the chromatin boundary protein Tbf1.

**GRAPHICAL ABSTRACT:** 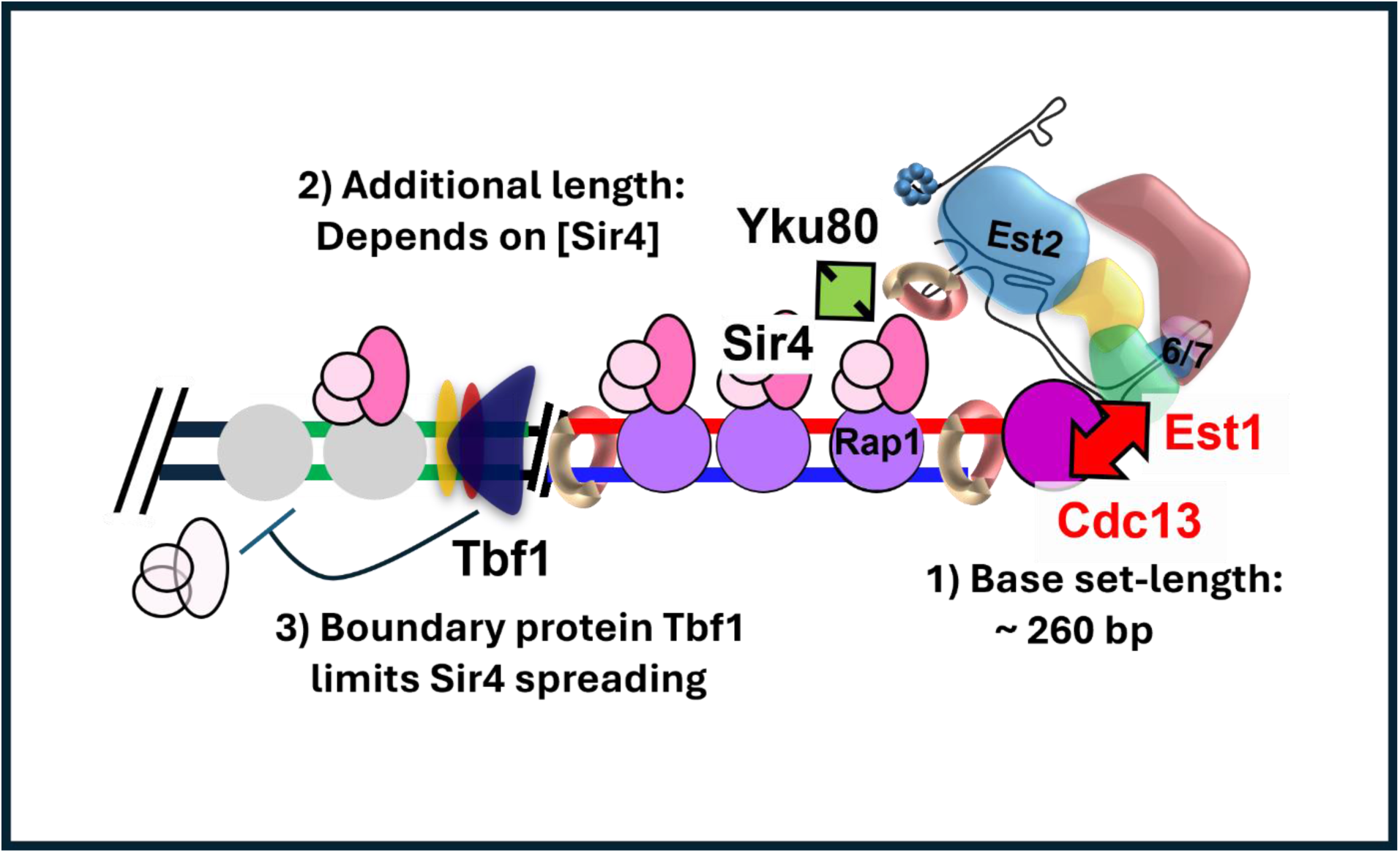

## INTRODUCTION

Telomeric repeat DNA and associated proteins are required for maintaining genome stability ^1^. In general, telomeric DNA is composed of short, guanine-rich repeats that render its replication particularly challenging ^2^. The so-called end-replication problem will lead to telomere shortening at every cell division ^3–5^, and replication fork difficulties can lead to abrupt losses of a large part of the repeats ^2^. Eventually, depending on the initial set-length of the telomeric repeat tracts, telomeric DNA becomes too short to assume telomeric functions and cells enter a permanent cell cycle arrest called cellular senescence ^6,7^. This telomere-length derived limitation of cell divisions on the one hand directly contributes to age-related degenerative disease, and on the other hand also restricts cells from undergoing uncontrolled cell divisions that may lead to cancer ^8,9^. Therefore, telomere equilibrium set-length is a key parameter that will affect the timing of onset of senescence and needs to be strictly regulated for optimal organismal function.

This telomere set-length is thought to be determined very early in life. Young humans have a set-length of about 10-12 kb of telomeric repeats ^10^. Currently, it is thought that very few, perhaps less than five, extremely short dysfunctional telomeres are sufficient to engage telomere-mediated cellular senescence ^11–13^. If the cellular DNA-damage checkpoint machinery is compromised, senescence can be bypassed, and most cells die in replicative crisis ^8,9^. However, some cells can pass crisis and advance to malignant transformation ^8^. Given that it takes only few dysfunctional telomeres to initiate the events leading to senescence, it may very well matter which are the causative telomeres for the process.

We know very little about how the initial set-length for telomeric repeat DNA is determined. Furthermore, while there is evidence that individual human ^14,15^ and yeast ^16,17^ telomeres can have telomere-specific set-lengths, mechanisms for how specific telomeres in the same cell could have significantly differing set-lengths remained unknown. Specifically, at least one budding yeast telomere, the telomere on chromosome end IIIL (TEL03L), has a set-length that deviates dramatically from those of all other telomeres ^18^. Indeed, in wild-type cells, the telomere set-length of TEL03L is about double the size of all other telomeres. Therefore, we investigated the mechanisms that lead to such a markedly different set-length for TEL03L. Significantly, our results show that an increased level of Sir4 at TEL03L is attracting telomerase via a Sir4-Yku80 interaction which links the TEL03L telomere to the telomerase RNA. This mechanism leads to an increased activity of telomerase at telomere TEL03L and delineates how the heterochromatin like domain on TEL03L leads to a telomere-specific telomerase recruitment mode that affects the telomere set-length determination exclusively in *cis*. Sir4 presence on telomeres is limited by heterochromatin boundary proteins such as Tbf1. We thus generated a mutation in Tbf1 that allows increased Sir4 presence, and in cells harbouring this new tbf1-453 allele, telomere set-length significantly increased on all telomeres. Therefore, our work on telomere TEL03L overall defines a molecular mechanism for telomere-specific set-length regulation. Extrapolating this new concept to human cells, such mechanisms could determine which telomeres have a higher or lower probability to be the first to become dysfunctional, leading to the initiation of senescence.

## RESULTS

### Extremely long telomere set-length on TEL03L is dependent on the Sir-proteins and the silent mating type locus HML

In budding yeast, either of two types of subtelomeric sequences, called Y’- or X-elements, occur directly adjacent to the terminal telomeric repeats ^19^. Accordingly, telomeres are classified as either X-telomeres or Y’-telomeres, each accounting for about half of all 32 telomeres in common lab-strains; 15 and 17 respectively in the strains used here. X-elements are not highly conserved, and many have enough unique sequences to be distinguished for single telomere analyses. During work on telomeric chromatin of such individual X-telomeres we found that the telomeric repeat tract on the X-telomere 3L consistently was much longer (510 ± 55 bp) than the previously described average of 300 ± 75 bp ^19^ (Suppl. Fig. 1a). This exceptional set-length for TEL03L was determined via TeloPCR (see Methods for details; Suppl. Fig. 1a) or by standard Southern blotting of genomic DNA digested with an appropriate restriction enzyme (PvuII, Fig. 1a, c; lanes WT). DNA from the same WT strains analyzed using the more common XhoI restriction digest and Southern blotting using a telomeric repeat probe did yield the canonical expected tract sizes on Y’-telomeres or on X-telomeres TEL11L and TEL06R (Fig. 1a-d; lanes WT). The fact that TEL03L is exceptionally long had been reported when this telomere was first cloned ^16^, and more recent data show that this occurs in most WT yeast strains from around the globe, including all lab strains ^18^. We also performed pan-telomere analysis by chromosome end-to-end sequencing using nanopore sequencing ^17^. Individual telomeric tract lengths are documented in violin plots for each telomere (Suppl. Fig. 1b WT, light blue) and compared to the average size of all telomeres (Suppl. Fig. 1b, stippled line). Analysis of the means of each telomere versus the grand mean of all telomeres shows that TEL03L, and to a lesser extent TEL08L, TEL08R, TEL09R and TEL11L were significantly longer, while TEL10R, TEL11R, TEL14L, and TEL14R appeared somewhat shorter than the grand mean of all telomeres (Suppl. Fig. 1c). Given that TEL03L was already confirmed to have an extremely long set-length ^16,18^, we concentrated our efforts on this telomere.

**Figure 1:**
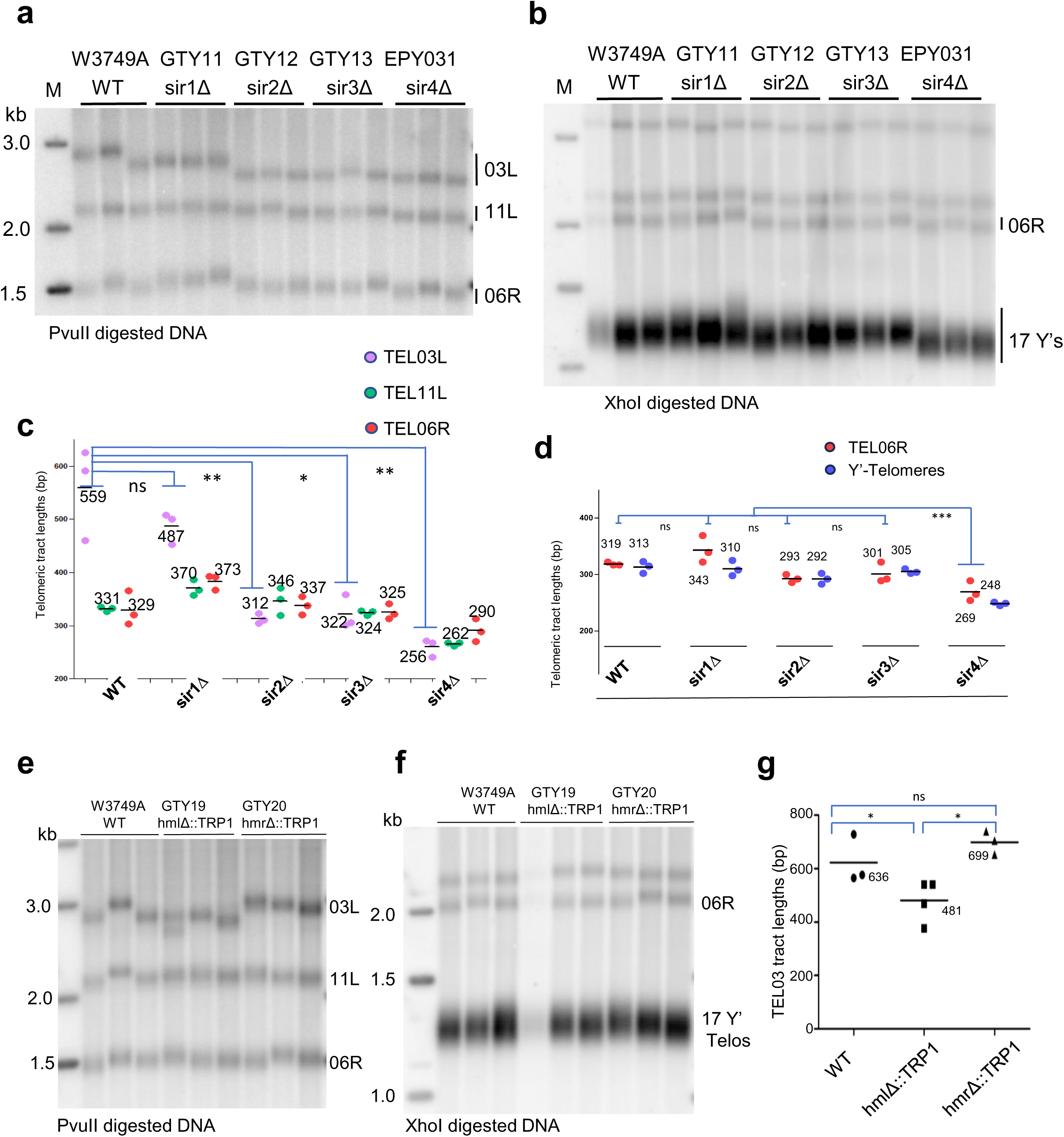
The long telomere set length on TEL03L is dependent on the SIR-proteins. ***a***. Southern blot of DNA derived from W3749A WT, *sir1Δ* (GTY11), *sir2Δ* (GTY12), *sir3Δ* (GTY13) and *sir4Δ* (EPY031) cells, digested with PvuII and probed with a telomeric repeat fragment. ***b***. Southern blot of the same DNA as in ***a***, but digested with the XhoI enzyme. ***c***. Quantified telomeric tract lengths from *a* of TEL03L, TEL11L and TEL06R are plotted with respect of the genotype (ns for difference not significant, ∗ for p < 0.05 and ∗∗ for p < 0.01, ∗∗∗ for p < 0.001, t-test). ***d.*** Quantification as in ***c.***, but for the blot in ***b.*** with the Y’-telomeres and TEL06R. ***e.*** Southern blot of DNA derived from W3749A WT, *hmlΔ::TRP1* (GTY19), and *hmrΔ::TRP1* cells, digested with PvuII and probed with a telomeric repeat fragment. ***f***. Southern blot of the same DNA as in ***e*,** but digested with the XhoI enzyme. ***g.*** Quantification of the telomeric repeat tract lengths on TEL03L in WT, *hmlΔ::TRP1,* and *hmrΔ::TRP1* cells. ∗ for p < 0.05, t-test.

Unique features of budding yeast chromosome III include a locus that determines haploid mating behavior (*MATa* or *MATα*). In addition, there are silenced copies of those mating type specifying genes near the telomeres of chromosome III: *HMR*, about 20 kb from telomere TEL03R ^20^; and *HML* about 10 kb from TEL03L ^16^. Silencing of those *HM*-loci depends on the association of a set of Sir-proteins (Sir1, Sir2, Sir3 and Sir4) ^21^. In fact, transcriptional silencing can also be observed near telomeres ^22^, and this repression is dependent on the Sir2, Sir3 and Sir4 proteins, but not on Sir1 ^23^. Focussing on telomere TEL03L on chromosome III, we therefore assessed whether the very long telomere set-lengths observed above are dependent on the presence of the SIR-genes (Fig. 1a-d; Suppl. Fig. 1b). In fact, deletion of any of the *SIR2, SIR3* or *SIR4* genes shortened the set-length telomere phenotype of TEL03L specifically such that the set-length of this telomere now became close to the median of all telomeres in the cell (Fig. 1a, c; Suppl. Fig. 1b). Such a Sir-gene specific deviation was not observed for any of the other telomeres on which the set-length deviated from the grand mean of all telomeres (Suppl. Fig. 1b). As reported previously ^24^, we also noticed that the deletion of the *SIR4* gene caused a slight reduction in the size of all telomeres, and remarkably, TEL03L also further adjusted to the slightly shorter overall set-lengths (Figure 1a-d). Specifically, the average telomere length in a *sir2Δ* strain measured 342±20 bp and that of TEL03L matched that size quite well 312±10 bp; in a *sir4Δ* strain, overall lengths were reduced to 276±21 bp and that of TEL03L to 256±14 bp (Figure 1a, c). Remarkably, in a *sir1Δ* strain, the overall telomere set-lengths were not significantly affected (Fig. 1c, d), but the length of TEL03L was reduced from around 559±87 bp to 487±30 bp (Fig. 1a, c). Given that the Sir1 protein exclusively is involved in silencing the HM loci, but not at telomeres, we surmised that Sir1 binding at the *HM*-loci may be required to affect the set-length telomere length in *cis*. We thus deleted *HML*, or *HMR* as a control, from the genome and discovered that deletion of *HML*, but not *HMR*, indeed also partly suppressed the long set-length phenotype on TEL03L (Fig. 1e-g). Notably, the set-length of the telomeric repeats on TEL03L in the *sir1Δ* strain (487±30 bp, Fig. 1c) was very similar to the size as in the *hmlΔ* strain (481±78, Fig. 1g). On the other hand, deletion of *HMR* did not affect any telomere set-lengths that we could measure (Fig. 1e-g). Further, neither of the deletions affected telomeric repeat lengths of X-telomeres TEL11L or TEL06R (Fig. 1e, f), and the overall length of all 17 Y’-telomeres also remained unaffected (Fig. 1e, f). We conclude that in WT cells, the set-length of TEL03L is dramatically longer than that of any other telomere and this extended set-length is dependent on the Sir2, 3, 4-proteins. Furthermore, the loss of Sir1 or the deletion of the *HML* locus exclusively caused TEL03L to shorten by about 100 bp and did not affect any other telomere set-lengths.

### Set-length telomere regulation on TEL03L depends on the telomerase pathway

We next wished to investigate how this very long equilibrium set-length at TEL03L was established and maintained. Telomere maintenance can be dependent on homologous recombination or on the telomerase enzyme ^19^. Examining telomere lengths by Southern blotting in strains lacking Rad52 and which can not maintain telomeres by recombination showed that the set-lengths of TEL03L were indistinguishable in WT vs *rad52Δ* strains (Fig. 2a). Therefore, we surmised that the TEL03L telomere was maintained by a telomerase dependent mechanism. In this case, the steady-state telomere length will be determined by an equilibrium between telomere shortening activities and telomerase-mediated telomere lengthening activities. To examine whether the length of TEL03L was due to reduced shortening activities, we compared the loss rates at various telomeres to that at TEL03L in the absence of telomerase. Previous data showed that in strains lacking an essential telomerase component, telomeres lose approximately 3-5 bp/generation ^25,26^. Given that we wanted to examine those losses immediately after telomerase loss, we fused Est3 and Est1, essential protein components of telomerase ^19^, to the FKBP12-rapamycin-binding domain of human TOR1 (FRB-domain) in the Anchor Away system ^27^. This setup allowed us by simple addition of rapamycin to rapidly deplete nuclear Est3/Est1 and examine telomeric sequence losses thereafter (Fig. 2b-e). At the start of the experiments, telomere lengths for Y’-telomeres as well as those for X-telomeres TEL11R and TEL06R were in the range of 300 ± 75 bp (Fig. 2b). However, TEL03L was much longer at approximately 480bp. We then induced Est3-FRB expulsion from the nucleus by addition of rapamycin and measured telomere length changes during the next ten days (corresponding to ∼95 population doublings (Fig. 2c, e). At the end, telomere loss rates were calculated as a function of population doublings for each X-telomere separately and the Y’-telomeres collectively (Fig. 2d). Although there is some variability in the data, the sequence loss rates at TEL03L were not significantly lower than those for the other telomeres analyzed in either the Est3-FRB or the Est1-FRB experiments (Fig. 2e). This result is consistent with previous results indicating that upon loss of telomerase, overall similar sequence loss rates are observed, irrespective of the starting telomere length ^28^. By inference, these data also strongly suggest that the extended set-length of TEL03L is due to a more efficient lengthening by telomerase.

**Figure 2:**
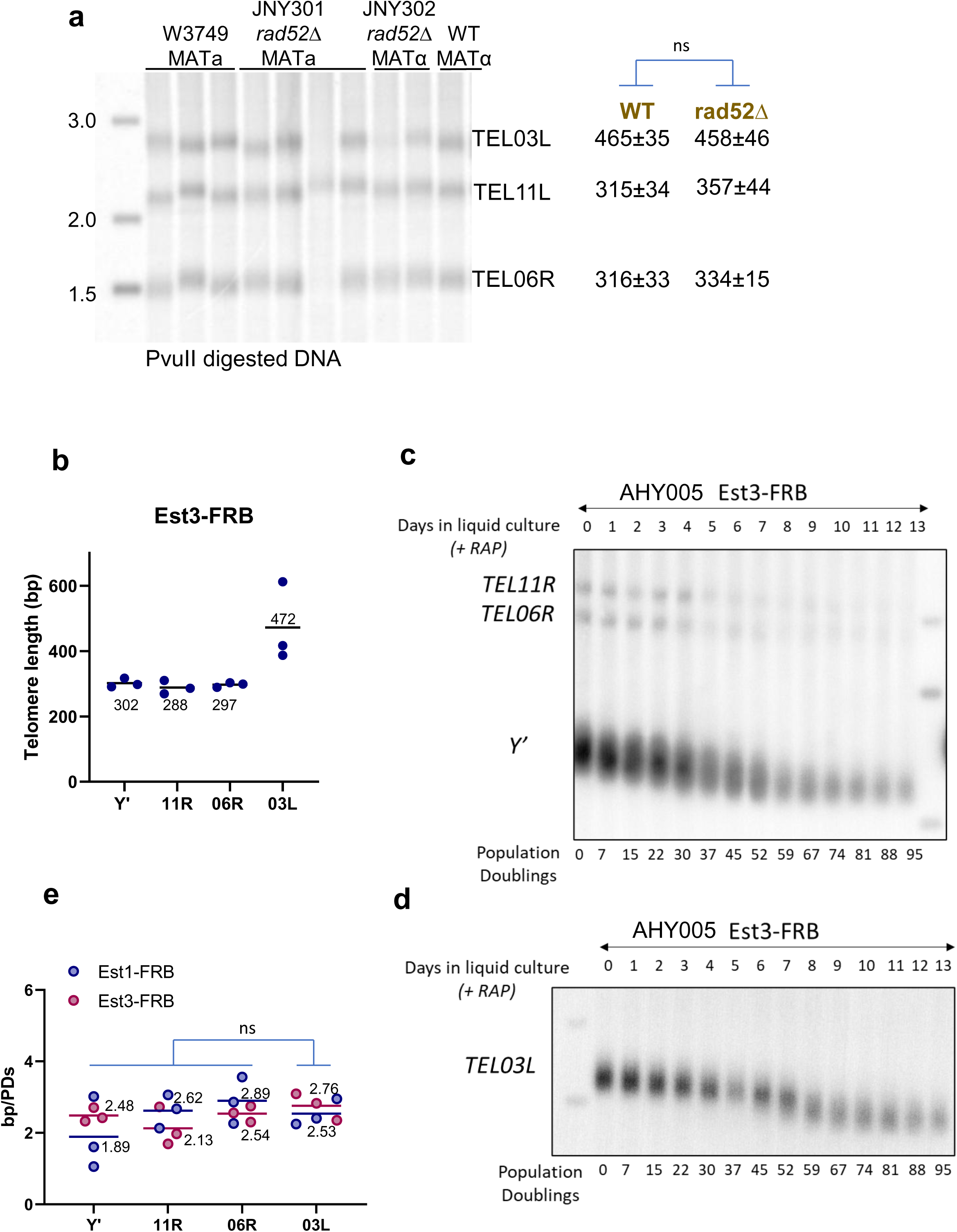
Long telomere set length regulation is dependent on the telomerase pathway. ***a***. Southern blot with DNA derived from three clones of W3749A, four clones of *rad52Δ MATa*, two clones of *rad52Δ MATα* and one clone of W3749α. DNA was digested with PvuII and probed with a telomeric repeat fragment. TEL03L, TEL11L and TEL06R telomeric repeat tract lengths are indicated on right. (ns for difference not significant, t-test). ***b.*** Initial telomeric tract lengths for 3 clones of Est3-FRB are shown for TEL03L, TEL06R, TEL11R and Y’-telomeres. ***c***. Changing telomeric tract lengths of TEL11R, TEL06R and Y’-telomeres during culture outgrowth (14 days or indicated population doublings) of Est3-FRB cells. ***d***. Same as ***c.*** but for TEL03L. ***e***. Calculated telomeric repeat loss rates (in bp/population doublings; bp/PDs) for Est1-FRB and Est3-FRB strains are tabled for TEL03L, TEL11R, TEL06R and Y’-telomeres. (ns for difference not significant, t-test).

### Long set-length regulation on TEL03L is contained in the distal 16.7 kb of subtelomeric DNA and is transferable

The extended set-length of the telomeric tracts on TEL03L occurs in many yeast strain backgrounds, including wild yeast strains collected in nature ^18^. Furthermore, the sequences on TEL03L, including the terminal repeats themselves, are highly conserved between strains. Therefore, the terminal sequence of TEL03L itself might have been at the base of the telomere-specific long set-length. We considered that either the TEL03L X-element or some special terminal repeat arrangement could cause the long set-length. To examine those two possibilities, we replaced the terminal TEL03L X-element with the X-element present on TEL01L (Fig. 3a, b; Suppl. Fig. 2a for details). As a control to examine the terminal repeat sequences only, the replacement procedure was also performed with the original TEL03L X-element that harbored only a very short telomeric repeat seed, followed by assessing the regrowth of the TEL03L in this situation (see Suppl. Fig. 2a). Remarkably, both situations allowed for a lengthening of the modified TEL03Ls to well beyond the average 300 bp size of telomeric repeat tracts in the cells: telomeric tract lengths as measured by Southern blotting were 420±40 bp for TEL03L-3L and 451±56 bp for TEL03L-1L (Fig. 3a, b). Similar results were obtained when the altered TEL03Ls were analyzed by TeloPCR (Suppl. Fig. 2b). The TEL03L before modification was at 466±65 bp. These data therefore show that the TEL01L X-element and a short telomeric repeat seed alone are sufficient to allow the generation of the long set-length on TEL03L. In addition, there is no special TEL03L-specific terminal repeat arrangement required for this overextension.

**Figure 3:**
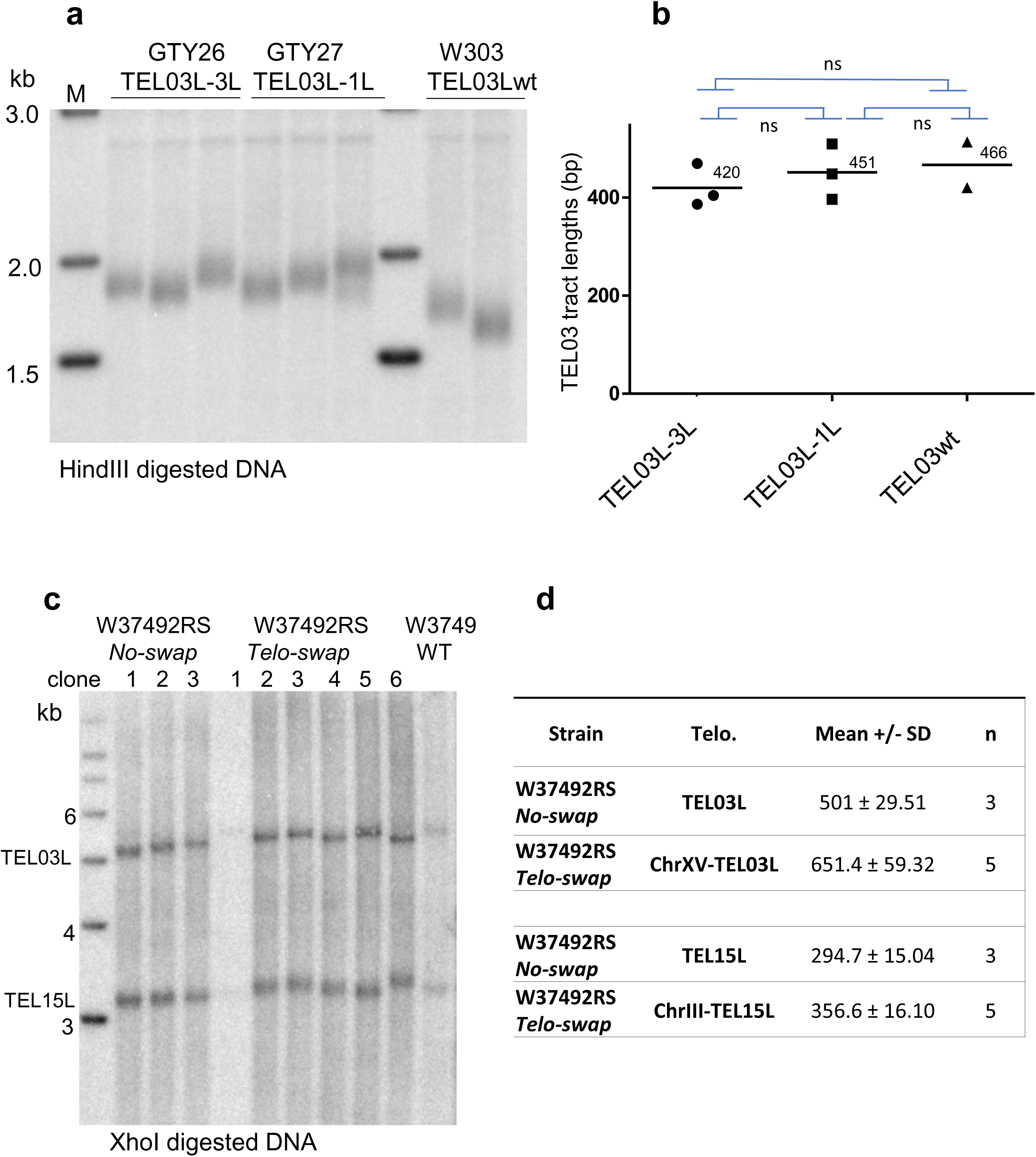
Telomere set length regulation can be moved to another telomere. ***a***. Southern blot and telomeric tract lengths of Tel03L-03L cells (GTY26), Tel03L-1L cells (GTY27) and WT cells with DNA digested with HindIII and probed with a TEL03L unique probe. ***b***. Quantified telomeric tract lengths (TEL03L) of the blot shown in *a*. (ns for difference not significant, t-test). ***c***. Southern blot of DNA derived from 3 independent clones of W37492RS No-swap cells, and from 6 independent clones in which the distal parts of TEL03L and TEL15L were swapped (W37492RS Telo-swap; see Suppl. Fig. 3 for details), digested with the XhoI restriction enzyme and probed with two unique probes for TEL03L and TEL15L. Lane far right: DNA from W3749 WT cells before RS site integrations. ***d***. Quantified telomeric tract lengths of TEL03L on chromosome III (TEL03L) or on chromosome XV (ChrXV-TEL03L) as well as TEL15L on chromosome XV (TEL15L) and on chromosome III (ChrIII-TEL15L).

Considering the partial suppression of set-length regulation on TEL03L observed in *sir1Δ* and *hmlΔ* strains (Fig. 1), the above data suggested that the genomic area from *HML* to the terminal repeat tracts may be required to specify the full extended set length on TEL03L in *cis*. If this was the case, this area alone should be sufficient to transfer an extended set-length to another telomere. We tested this hypothesis by swapping the entire distal telomeric TEL03L region (∼ 16.7 kb including *HML*) with the distal 13.5 kb telomeric region of TEL15L on chromosome XV (Fig. 3c, d; see schematic in Suppl. Fig. 3a for details). The telomere swap via the inserted RS recombination sites was verified by locus-specific PCR (Suppl. Fig. 3b) and by hybridization of pulsed field gel (CHEF) Southern blots (Suppl. Fig. 3c). Telomeric tract lengths on the original or transposed telomeres TEL03L or TEL15L was then assessed by Southern blotting (Fig. 3c, d). The displaced TEL03L on chromosome XV (ChrXV-TEL03L) indeed was sufficient to maintain a very long telomere set-length (650 ± 60 bp), while the TEL15L telomeric area on chromosome III (ChrIII-TEL15L) remained close to the average length of all other telomeres (356 ± 16 bp, Fig. 3d). These data thus show that the distal telomeric region of chromosome III enforces an extended set-telomere length exclusively on this telomere in *cis*, independently from the identity of the rest of the chromosome.

### Telomerase recruitment via Sir4-yKu is required for long set-length on TEL03L

The above data suggest that the extension activity of telomerase is more active on TEL03L than on other telomeres. Our hypothesis was that telomerase-mediated extension and hence telomerase recruitment somehow was more efficient on TEL03L. Telomerase recruitment in yeast depends on an essential interaction between the Est1 component on telomerase and the telomere-bound Cdc13 protein ^19^. However, the yeast yKu heterodimer, composed of Yku70/Yku80, interacts with the telomerase RNA Tlc1 ^29^ and is involved in non-essential alternative recruitment of telomerase. Specifically, Yku80 interacts with Sir4, which in turn can interact with the telomere-bound Rap1 protein. This Tlc1-Yku70/Yku80-Sir4-Rap1 axis provides for accessory telomerase recruitment that lengthens all telomeres by about 50 bp ^30,31^. Given the above, the long set-length on TEL03L could be due to an enhancement of this recruitment mode. This hypothesis predicts that in strains harboring any mutations that disrupt this axis, all telomeres would be equally dependent on the Est1-Cdc13 recruitment and have the same set-lengths. We therefore introduced alleles that interrupt the axis and compared global telomere set-lengths to the one on TEL03L. The telomerase RNA allele *tlc1-Δ48* does not interact with yeast yKu ^29^ and the *yku80-L140A* allele of *YKU80* can not interact with Sir4 ^32^. The average telomere length of all telomeres decreased slightly by about 50 bp in strains with the *tlc1-Δ48*, *yku80-L140A* or the *sir4Δ* alleles (Fig. 4a, b; Suppl. Fig. 4a, b), consistent with previous data ^29,32^. Remarkably, in all three strains, the set-length of TEL03L also was reduced to the same lengths as the mean of all telomeres (Fig. 4b; Suppl. Fig. 4a). These results thus show that the long telomere set-length of TEL03L is dependent on the Tlc1-Yku70/Yku80-Sir4 telomerase recruitment axis, which therefore mediates the telomere-specific level of telomere-telomerase interactions.

**Figure 4:**
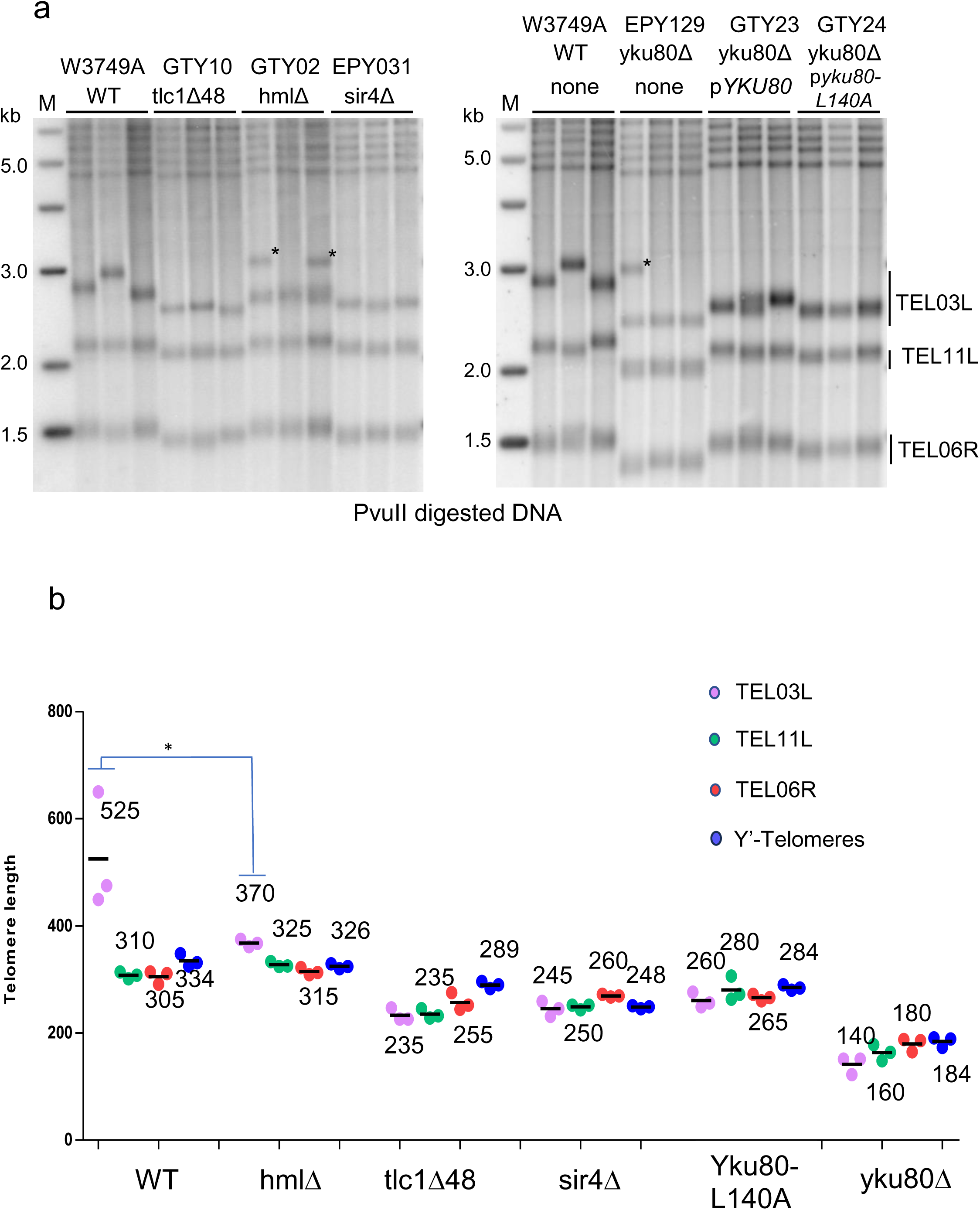
Long set-length on TEL03L depends on telomerase recruitment via the Sir4-yKu-Tlc1 axis. ***a.*** Left: Southern blot of DNA derived from W3749A WT cells, cells harboring the *tlc1Δ48* allele (GTY10), cells harboring the *hmlΔ::URA3* allele (GTY02), or cells with a *sir4Δ* allele (EPY031). DNA was digested with PvuII and hybridized to a telomeric repeat fragment. Right: Southern blot as on left, but with W3749A WT cells, cells with a *yku80Δ* allele (EPY129), EPY129 cells in which the *yku80Δ* allele is complemented by a plasmid-borne WT *YKU80* (indicated as p*YKU80*; strain GTY23) and EPY129 cells in which the *yku80Δ* allele is complemented by a plasmid-borne *yku80-L140A* allele (indicated as p*yku80-L140A*; strain GTY24). Bands labeled with an * are new recombinant X-telomere bands, but not TEL03L (see Supplmental Fig. 4a). ***b.*** Quantified telomeric tract lengths of TEL03L, TEL11L, TEL06R, and Y’-telomeres are plotted on the graph vs the indicated genotypes (ns for not significant, ∗ for p < 0.05, t-test). Southern blots for the data are in Fig. 4a, Suppl. Fig. 4a, b)

### A mutation in Tbf1 increases telomeric Sir4 association and globally increases telomere set-lengths

There is evidence that on TEL03L, the Sir2/3/4-proteins spread from the distal telomeric X-element and are associated with the entire 10 kb sequence between the *HML* locus and TEL03L ^33,34^. This explains a significantly increased presence of Sir4 on this telomeric locus but this Sir-protein spreading on TEL03L is exceptional on telomeres. On virtually all other natural telomeres, the Sir-proteins do not spread much beyond the telomeric repeat DNA on Y’-telomeres, or beyond the proximal end of the X-elements ^33,34^. It is therefore stipulated that except for the proximal end of the X-element on TEL03L, there is a strong chromatin boundary that inhibits spreading beyond the transition sites ^33–35^. Given that the telomeric repeat tract distal of the X-element of TEL01L also becomes very long if that X-element is placed onto the end of telomere IIIL (Fig. 3), we hypothesized that Sir-protein coverage from TEL03L to the *HML* locus may be due to telomere-independent association of Sir-proteins at the *HML* locus and spreading towards the telomere, an effect that could weaken this boundary element in *cis*.

If indeed a weakened boundary element at TEL03L caused the long set-length on TEL03L, mutations in boundary element proteins, such as Tbf1 ^35^, may weaken the boundary effect in general, allowing increased Sir4 association on all telomeres and leading to overall telomere lengthening. Boundary element proteins such as Tbf1, Reb1, or Abf1 are also important general transcription factors and essential in yeast ^19^. Therefore, we screened for a new allele of *TBF1* that still supports cell viability but in which the boundary function may be affected. The *tbf1-453* allele fulfills these requirements: it contains a single amino acid change (Q453H) and cells harbouring this allele grow slowly but are viable at 30°C. In these cells, no major change in their transcriptomic profiles was detectable (Suppl. Fig. 5a, b, c). Moreover, in cells with the *tbf1-453* allele, the abundance of the Sir4 protein significantly increases on telomere adjacent sites (1.5 kb to 3.1 kb from the transition of subtelomeric to the respective X-elements; Suppl. Fig. 5d). Finally, in these cells, all analyzed telomeres had extended set-lengths, including X-telomeres TEL06R, TEL11L, TEL11R, TEL03L, and the Y’-telomeres (Fig. 5a, b, Fig. 6). Moreover, these new telomere set-lengths are specific to the *tbf1-453* allele and are not observed in cells lacking another telomeric chromatin associated protein such as Rpd3 (Fig. 6a, b). Deleting any of the *SIR2, SIR3 or SIR4* genes in cells with the *tbf1-453* allele however is epistatic and reduces the elongated set-length of the telomeres (Fig. 5b), as predicted if the over elongation of telomeres in the *tbf1-453* cells is dependent on Sir-protein association on telomeres.

**Figure 5:**
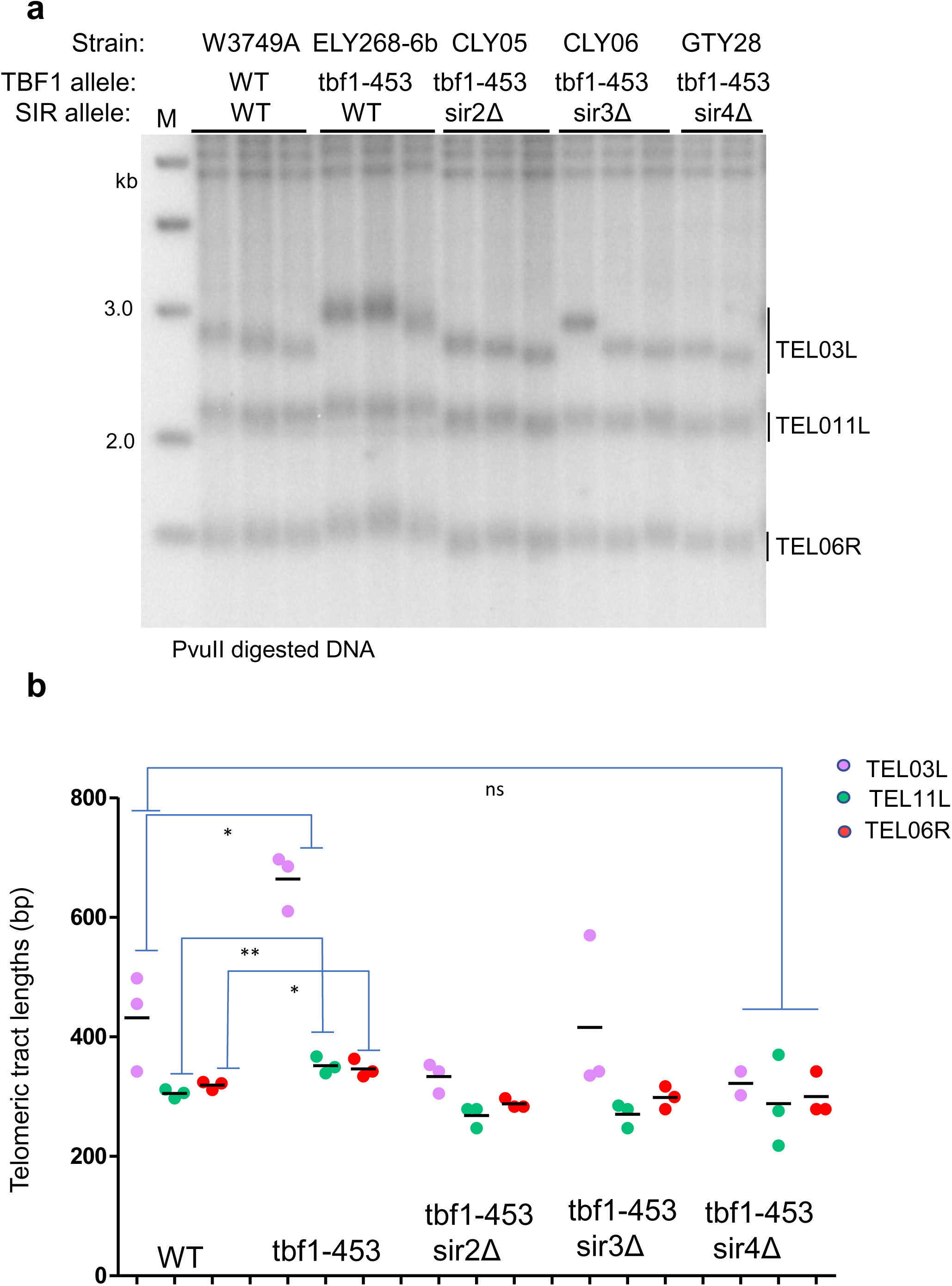
A boundary element mutation causes Sir-dependent telomere elongation. ***a.*** Southern blot of DNA from W3749A WT cells, cells with the *tbf1-453* allele (ELY268-6b), cells with both the *tbf1-453 and sir2Δ* alleles (CLY05), cells with both the *tbf1-453 and sir3Δ* alleles (CLY06), cells with both the *tbf1-453 and sir4Δ* alleles (GTY28). DNAs were digested with PvuII and probed with a telomeric repeat fragment. ***b.*** The telomeric tract lengths of TEL03L, TEL11L andTEL06R from the gel in *a* were quantified and plotted on the graph vs the indicated genotype (ns for not significant, ∗ for p < 0.05 and ∗∗ for p < 0.01, t-test).

**Figure 6:**
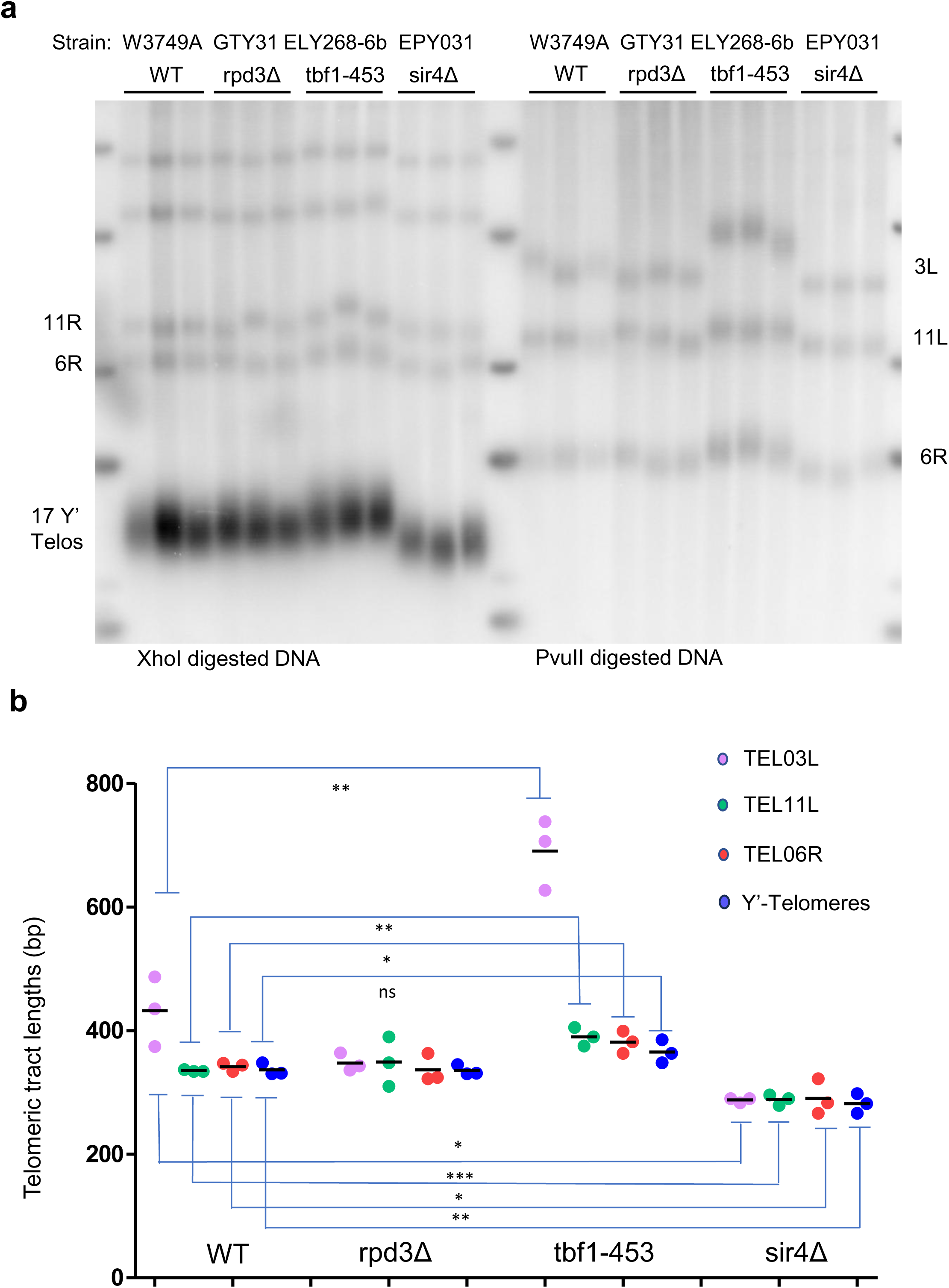
Rpd3 is not involved in telomere set-length regulation. **a.** Southern blot of DNA from W3749A WT cells, cells with the *rpd3Δ* allele (GTY31), cells with the *tbf1-453* allele (ELY268-6b), and cells with the *sir4Δ* allele (EPY031). The DNA was digested with XhoI (left part), or PvuII (right part) and hybridized to a telomeric repeat fragment. ***b.*** The quantified telomeric tract lengths of TEL03L, TEL11L and TEL06R vs the indicated genotypes are plotted (∗ for p < 0.05, ∗∗ for p < 0.01 and ∗∗∗ for p < 0.001, t-test).

## DISCUSSION

The analyses presented here show that on TEL03L in yeast, an increased presence of Sir-proteins on the terminal 10 kb of DNA is responsible for a very long equilibrium set-length of telomeric repeats specifically at this telomere. It is well established that the Sir-proteins associate with regions requiring transcriptional silencing, such as the *HML* and *HMR* loci ^21^. In addition, the Sir2/3/4-proteins, but not Sir1, also strongly associate with telomeres and subtelomeric X-elements ^33,34^. However, on all native telomeres except TEL03L, there is only very limited presence of the Sir2/3/4-proteins centromere proximal from the X-elements ^33,34^. The abrupt loss of Sir-association at the centromere proximal side of the X-elements or at the transitions between telomeric repeat DNA and the Y’-element closely parallels the loss of transcriptional repression in native telomere environments, an effect ascribed to boundary elements at these transition sites ^35,36^. Our data show that in an experimental setting with generally increased Sir-protein associations; in cells with the *tbf1-453* allele, all analyzed telomeres, including TEL03L, lengthen to a new set-length that is approximately 50 bp longer than in WT cells (Fig. 5, Fig. 6).

Next, our data show that the very long set-telomere length on TEL03L is dependent on the yKu-Sir4 protein interaction that bridges Sir4-bound telomeric loci to yKu-bound telomerase RNA (Fig. 4). It was reported before that for Y’-telomeres, this non-essential telomerase recruitment mode results in a 50 bp increase of the telomere set-length from about 260 bp, in the absence of the yKu-Sir4 axis, to about 300 bp in WT cells ^24,29^ (Fig. 4). However, on TEL03L, the set-length is increased from 260 bp to ≥ 500 bp (Fig. 1) in a telomerase-dependent manner (Fig. 2). The X-element on TEL03L is not required for the long set-length on TEL03L, because replacing this 03L-X with the one of TEL01L (01L-X) also results in a very long set-length on TEL03L (Fig. 3). By extension, the boundary element on the TEL03L-X does not appear to differ from other boundary elements. Interestingly, the presence of the *HML* locus does affect the set-length of TEL03L by a significant amount (Fig. 1e-g), but has no effect on the set-lengths of any other telomere. We ascribe this result to the strong association of the Sir-complex with the *HML* locus (discussed above). Consistent with this idea, the fully elongated set-length at TEL03L is dependent on the Sir1 protein and the presence of the *HML* locus, but the equilibrium set-lengths of other telomeres are not significantly influenced by their presence or absence (Fig. 1, ^23^). Finally, transposing the entire distal region of chromosome III onto chromosome XV leads to an extended telomere set-length on the transposed telomere, while the new telomeric locus on chromosome IIIL comprising the chromosome XV specific distal sequences remains in the 300-350 bp range, characteristic of many other telomeres (Fig. 3).

These results therefore detail the molecular requirements for how the very long set-length of TEL03L is achieved: an increased association of the Sir4 protein over 10 kb of telomere-adjacent chromatin yields increased Sir4-yKu mediated telomere-telomerase interactions. The latter allows for an increased probability of telomere elongation events in *cis* on this telomere, leading to a long set-length telomere. We surmise that the probability of extensions events still is inversely correlated with repeat-length, as previously proposed ^37^, but that on TEL03L, the probabilities are increased for longer telomeres. Given a constant shortening rate for all telomeres (Fig. 2), the telomere set-length will thus be increased at equilibrium. This mechanism also explains the previously puzzling observation that tethering of the Sir4 protein to a specific telomere will cause lengthening exclusively of that particular telomere in *cis* ^38^.

Other telomeres also deviate from the mean of all telomeres (Supp. Fig. 1 and 17), but so far, it remains unclear whether their regulation is based on the mechanism described here or additional mechanisms exist to regulate end-specific equilibrium telomere lengths.

Taken together, our results document a mechanism for how the equilibrium set-length of telomeric repeats can be determined by specific subtelomeric chromatin in *cis* and thus lead to a telomere-specific length homeostasis. Recent data indicates that in human cells, telomeres also have chromosome end-specific set lengths ^14,15^. For humans, telomere length is a critical and limiting parameter for the number of cell divisions that can occur. For example, very short telomeres lose their full capping function and are sensed by a telomere-specific DNA damage response that induces cellular senescence ^6,39^. Moreover, inappropriate telomere lengths are associated with cancer development. Chromosome end-specific length differences predict distinct telomeres may be the first to be sensed as dysfunctional, which will have important implications for aging and cancer.

## METHODS

### Strains and plasmids

All strains are derivatives of W3749 *MATa*, a *RAD5*-corrected W303 strain ^40^ unless otherwise noted. The genotypes of the strains used in this study are described in Supplemental Table 1.

To delete Sir genes, we constructed GTY11, 12, and 13. These strains were obtained by a one-step replacement of the *SIR1, SIR2* and *SIR3* genes with a KanMX6 cassette. To delete *RAD52* we constructed JNY301 and JNY302 by a one-step replacement of *RAD52* with a *LEU2* cassette in W303 *MATa* and *MATα* ^41^.

For the anchor away system we constructed AHY001 and AHY005 strains from KLY18 strain by C-terminal tagging of *EST1* and *EST3* genes respectively by one-step PCR-mediated technique. The FRB::KanMX6 cassettes used for C-terminal tagging of Est1 and Est3 were obtained from amplification of pLK2 plasmid with primers Est1_CterTag_F and Est1_CterTag_R and Est3_CterTag_F and Est3_CterTag_R (Supplemental Table 2).

For the TEL03L X-element swap we constructed GTY26 and GTY27. GTY26 was constructed by transforming W303 *MATa* ^41^ with a plasmid containing TEL03L subtelomeric region, Tel03L X-element and telomeric repeats with a *LEU2* flanked by RS site (T3TD plasmid digested with NsiI/Eco53kI). After, the *LEU2* selection marker was recombined out obtaining the final strain. GTY27 was constructed by transforming W303 *MATa* WT with a plasmid containing TEL03L subtelomeric region, TEL01L X-element and telomeric repeats with a *LEU2* flanked by RS sites (pAH01 digested with NsiI/Eco53kI) that was recombined out to obtain the final strain.

For the deletion of *HML* and *HMR* loci, we constructed GTY02, GTY19 and GTY20. GTY02 was obtained by transformation of W3749A with pJH2039 ^42^ digested with HindIII-HF. GTY19 and GTY20 were obtained by transformation of W3749A with PCR fragments (primers Hml-F and R and Hmr-F and R (Supplemental Table 2) from plasmids pGT02 and pGT03 respectively.

For the deletion of the Ku stem of Tlc1 in W3749A strain, we constructed GTY10. Plasmid pRS306-tlc1Δ48 digested with BsrGI was used for the integration and then, PCR was used to select correct recombination. For expression of Yku80 and Yku80-L140A, EPY129 strain was transformed with the pJP7c plasmid (*YKU80-Mycx2-Hisx10*, CEN, *TRP1*) and pJP7C-L140A ^43^.

To construct *tbf-453* (ELY268-6b), site-directed mutagenesis primers (Supplemental Table 2) were designed to introduce the desired amino acid substitution into the pRS304-TBF1::NatMX plasmid which was then digested to obtain the desired fragment for integrating *tbf1-453* allele in the TBF1 locus to create a heterozygous diploid. Then, the strain was microdissected to obtain haploid strains. CLY05, CLY06 and GTY28 were obtained by a one-step replacement of the *SIR2, SIR3* and *SIR4* genes with a KanMX6 cassette in *tbf1-453* strains. Strain GTY31 was obtained by a one-step replacement of the *RPD3* gene with a *TRP1* cassette.

To do the W37492RS Telo-swap (TEL03L-TEL15L telomere swap) we constructed EDY10. We started by constructing EDY06 (RS-URA3-RS was added 16.8 kb away from the telomeric repeats of TEL03L-W3749α) and EDY07 (RS-URS3-RS was added 13.5 kb away from the telomeric repeats of TEL15L-W3749a). The URA3 flanked by RS sites was recombined out to obtain the final strains. These strains were mated, recombination was induced, and the strains were finally microdissected to obtain haploid strains.

Strains containing the *tbf1-453* and *sir4-myc* alleles were constructed by mating W3749 *tbf1-453* with a *SIR4-Myc* W303 (JC3433 W303 *MATa* with *SIR4-13Myc::KanMX*) ^44^. Then strains were microdissected to obtain haploid strains.

Yeast strains obtained were verified by PCR followed by sequencing or Southern blot. Unless specified differently, in all experiments, yeast cells were grown at 30°C with constant agitation in standard conditions (YEP or YC media with appropriate carbon sources).

Plasmids are described in Supplemental Table 3 and all plasmids used were verified by sequencing.

T3TD was obtained by Gibson Assembly to be able to insert the subtelomere of TEL03L, *LEU2* flanked by R-sites and the X-element of TEL03L into the NsiI-NotI sites of pRS303. Then, telomeric repeats were inserted into the NruI-NotI sites of the new plasmid.

pAH01 was obtained by replacing X-element of TEL03L with X-element of TEL01L in T3TD (NcoI-NotI sites).

pGT01 and pGT02 were constructed by Gibson assembly by adding HML or HMR flanking sequences with the *TRP1* marker to a pRS303 plasmid.

pEP21 was constructed by ligating pJP7C (NotI digested) with the NotI digested fragment of pB3 ^45^.

### DNA isolation and Southern blotting

DNA was extracted using a standard phenol–chloroform procedure ^46^. An appropriate quantity of digested DNA, ranging from 0.5 µg to 1 µg, was separated on 0.7% TBE (1 x) agarose gels, transferred onto a BrightStar™-Plus Positively Charged Nylon Membrane (Invitrogen) and hybridized to specific 32^P^-labelled radioactive probes. Specific probes (TEL03L, TEL15L) to one chromosomal end were obtained by genomic DNA PCR-based amplification using specific primers followed by amplicon purification on gel and random priming labeling procedure ^47^. Sequences of primers used to amplify a specific chromosomal end are depicted in Supplemental Table 2. As a telomeric repeat probe, a 300-bp fragment containing 280-bp TG repeats derived from pYLPV was radiolabelled using a random priming labeling procedure ^48^. Data were visualized with a Typhoon FLA9000 apparatus. When necessary, membranes were stripped by boiling in 0.1% SDS and left at room temperature for 30 min and rehybridized with other radioactive probes. Data analysis GelAnalyzer software was used to determine the size of each detectable fragment.

Pulsed field gels were run using a BioRad CHEF-DR II system. Full-length chromosomal DNA was prepared according to the manufacturer’s instructions. DNA was subjected to electrophoresis at 14°C in a 1% agarose gel in 0.5 x TBE for 24 hours at 200 volts with switch times ramping from 60 seconds to 120 seconds.

Following electrophoresis, the gel was stained with 1 µg/ml ethidium bromide and photographed, then the chromosomal DNA was fragmented by exposure to 60 mJoules of 254 nm irradiation. The DNA was transferred to a Nytran SuPerCharge membrane using a Whatman Turboblotter. Transfer was performed according to the alkaline method described by the manufacturer.

### TeloPCR

We modified the protocol based on Zubko *et al.* 2016 ^49^. For the terminal transferase-mediated tailing of genomic DNA, we used 60 ng of genomic DNA in a volume of 10 µL and the DNA was denatured at 95°C for 5 min in a thermal cycler and cooled on ice. Then, 40 µL of transferase reaction mix was added to make total volume 50 µl. The transferase reaction mix contained 1X transferase buffer, 0.25 mM CoCl_2_, 0.2 mM dCTP and 2 U of terminal transferase (New England Biolabs). The mix was incubated at 37°C for 30 min, and then the enzyme was heat inactivated at 70°C for 10 min at. After heat inactivation, the tailing reactions were used in PCR.

PCRs for amplification of chromosomal termini were performed on tailed DNA in 20 µL using GoTaq® Long PCR master Mix (Promega). PCR mix included 10 µL of GoTaq® Long PCR master Mix, 0.5 M of a subtelomeric specific region, 1 M of a G tail primer (Supplemental Table 2) and 1 µL of C-tailed genomic DNA. PCR conditions were the following:

95 °C - 2 min, and then 40 cycles at 95 °C - 20 s, 65 °C - 30 s, 65 °C - 2 min with final extension at 72 °C for 10 min. PCR reactions were carried out in 0.2 mL tubes (VWR) using Bio Rad T-100 Thermal Cycler. After PCR a 1.2% TBE 0.5X gel was run overnight at 40 volts.

### Nanopore sequencing

High molecular weight genomic DNA extraction from BY4741 WT ^50^ and BY4741 *sir3Δ* cells and DNA molecular end tagging were done according to Sholes *et al*. 2021 ^17^.

Nanopore library preparation of 1µg input DNA with a molecular tag was performed using the Native barcoding genomic DNA kit (Oxford Nanopore Technologies EXP-NBD104 and SQK-LSK109). Samples were run on a MinION flow cell (v.9.4.1) for 24 to 72 hours and were operated using the MinKNOW software (v.19.2.2). Only reads with barcodes on both ends were selected to pass.

To analyze individual telomere lengths from nanopore sequencing, software has been developed and the code is hosted on GitHub (https://github.com/jflucier/E2EAssembler). The first step allows genome assembly from Nanopore sequencing data for each strain. We divided the sequencing reads into different parts, each with a total genome coverage of approximately 60X. We used Canu assembly software to individually assemble each different part. The produced assemblies were then merged ^51^. The second step consists of the annotation of subtelomeres, telomeric repeats, and telotag on the assembled genome. The third step consists of the extraction of length value of telomeric tracts for each read from the different chromosome ends. Data representation was done using R software.

### Anchor away system

We modified the protocol based on Haruki *et al*. 2008 ^27^. Senescence assays of anchor-away strains have been performed with 2 WT clones, 3 independent clones of EST1-FRB and 3 clones of EST3-FRB. YEPD supplemented with rapamycin at 1 μg/ml final (YEPD + Rapamycin) was freshly made daily. After an overnight pre-culture in YEPD, cells were diluted at 0.04×10^7^ cells/ml in YEPD + Rapamycin. After 24 hours, OD_660_ was measured, and cells were diluted to 0.04×10^7^ cells/ml in YEPD + Rapamycin. Population doublings were calculated using OD_660_ measurements converted to cells/ml using a standardized chart. The process was repeated each day for 14 days and for each day, the remaining cells were pelleted, and DNA was extracted.

### Chromatin immunoprecipitation and qPCR

Chromatin immunoprecipitation experiments were performed as previously described ^52^ with minor modifications. Briefly, 100mL cultures per ChIP were grown at 30 °C to OD_600nm_ 0.5–0.8. Cells were cross-linked with 1% formaldehyde for 10 min. After stopping the cross-link reaction and washes, cells were lysed in 500 μL ChIP lysis buffer (50 mM HEPES–KOH pH 7.5, 140 mM NaCl, 1 mM EDTA pH 8.0, 1% Triton X-100, 0.1% Na-deoxycholate, 1 mM PMSF) with glass beads using a FastPrep (MP Biomedicals). Following sonication, 500 μL of whole cell extract (WCE) was incubated overnight at 4°C with 2 μg anti-*c*-myc antibodies (mouse monoclonal, clone 9E10, Roche) for immunoprecipitation of myc-tagged proteins. The next day, 50 μL equilibrated Pierce protein A/G magnetic beads (Thermo Fisher Scientific) were added for a 2 hour incubation at 4 °C. The purified immunoprecipitated and input DNAs were diluted and subjected to qPCR using appropriate primers (Supplementary Table 2) and PerfeCTa SYBR Green SuperMix (Quantabio, 95054-02K).

The percent input method was used to analyze qPCR signals. Briefly, the input signal was adjusted to 100% using the formula: Ct Input - log2 of dilution factor. The ΔCq was calculated using: Ct IP - adjusted Ct input. The percent input was then calculated using the formula:

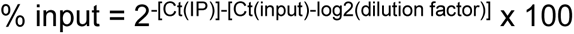

qPCR reactions were performed in technical triplicates for 3 independent biological cultures. The mean, standard deviation and statistics for independent biological replicates are reported

### Total RNA sequencing

For total RNA extractions, 10 mL of yeast cells were grown to approximately OD_600_ 0.8, washed, resuspended in 500 µL LETS buffer and lysed by glass beads. Samples were extracted twice with phenol/chloroform/isoamyl alcohol (25:24:1) and once with chloroform/isoamyl alcohol (24:1). RNA was precipitated with 30 µL 3M NaOAc and 1 ml cold 100% ethanol. Samples were washed with 70% EtOH, dried, and resuspended in 20 µL of nuclease-free water and quantified. RNA integrity was assessed with an Agilent 2100 Bioanalyzer (Agilent Technologies). Libraries were quantified and subjected to RNA-sequencing by Illumina PE100bp Hiseq 4000.

### Statistics and reproducibility

GraphPad prism was used to perform all statistical analyses. Statistical differences of mean values of measurements were determined by two-tailed unpaired Student t test. Significance was denoted as ns for not significant, ∗ for p < 0.05, ∗∗ for p < 0.01, ∗∗∗ for p < 0.001.

## DATA AVAILABILITY

- The complete numerical values for all telomere length determinations by Southern blotting reported in this paper will be shared by the lead contact upon request. The dataset on the whole genome raw sequencing reads generated in this study has been submitted to the NCBI BioProject database under accession number PRJNA1088003. The Illumina RNA-seq data have been submitted to the NCBI BioProject database under accession number PRJNA1099244.
- This original code/software for determining individual telomere lengths from nanopore sequencing, E2EAssembler, is hosted on GitHub and publicly available (https://github.com/jflucier/E2EAssembler).
- Any additional information required to reanalyze the data reported in this paper is available from the lead contact upon request.

## AUTHOR CONTRIBUTIONS

GMT and EP performed all biochemical and *in vivo* experiments; EB, EDL and LB contributed strain constructions, ChIP and RNA seq experiments; J-FL wrote scripts for nanopore sequencing analyses; DSD performed the pulsed field gels; RJW study conception, design and financing. RJW wrote and edited the manuscript with input of all co-authors.

## ACKNOWLEDGEMENTS

We thank the Wellinger lab for constructive discussions and A. Krallis, A. Hewitt, and C. Levasseur for experimental help during the early phase of the project and with the anchor away experiments. We also thank the Brewer/Raghuraman lab for tools used in the anchor away system. Finally, we thank S.L. Sholes and C. Greider for generously providing Nanopore sequencing and their input in the analyses. Work for this manuscript was supported by a grant from the Canadian Institutes for Health Research (CIHR; foundation grant FDN154315); funds from the Canadian Research Chair on Telomere Biology to RJW; a post-doctoral fellowships to GMT from the Sherbrooke Center for Aging Research (CdRV); and a post-doctoral fellowship to LB from the Fonds de Recherche du Québec – Santé (FRQS). DSD was supported by NIH grant R35 GM152165.

