## Supplemental Information for "A mechanism for telomere-specific telomere length regulation"

**Supplemental Figures 1-5**

**Supplemental Tables 1-3**

### Supplemental Figure 1

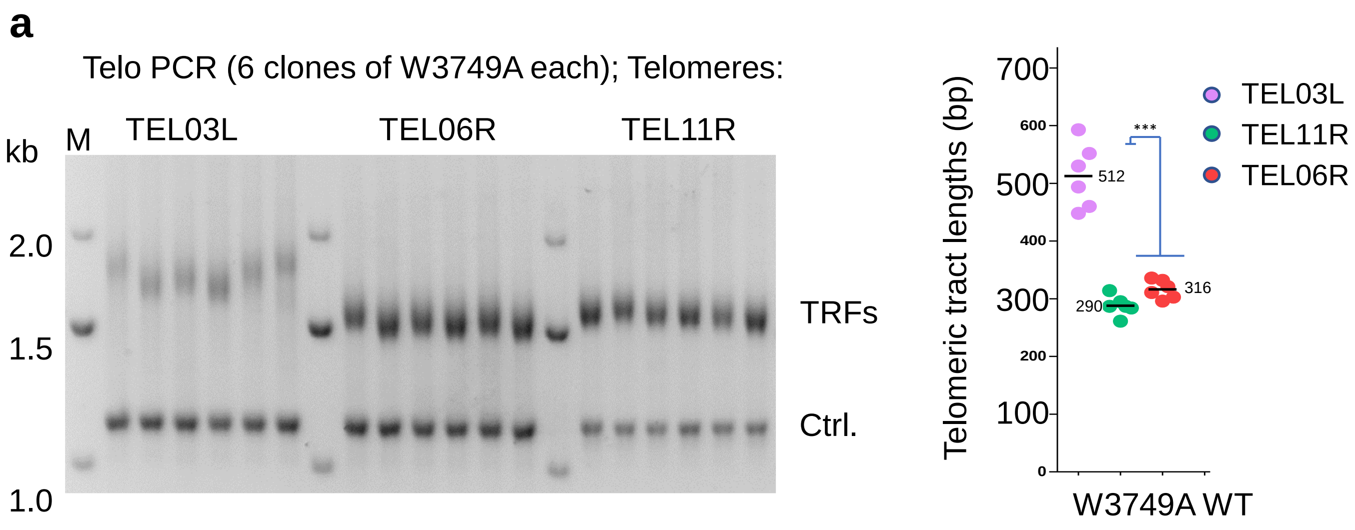


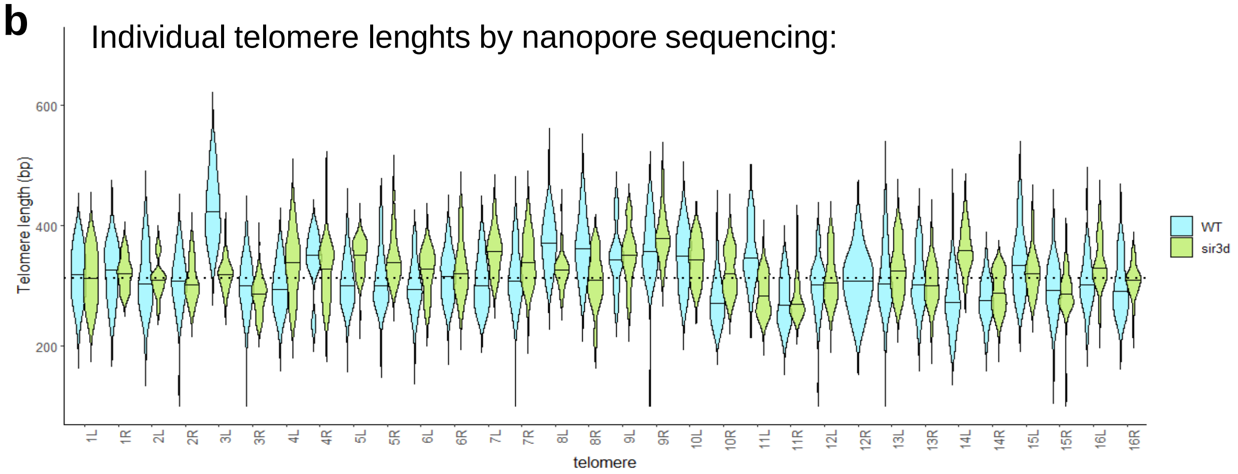


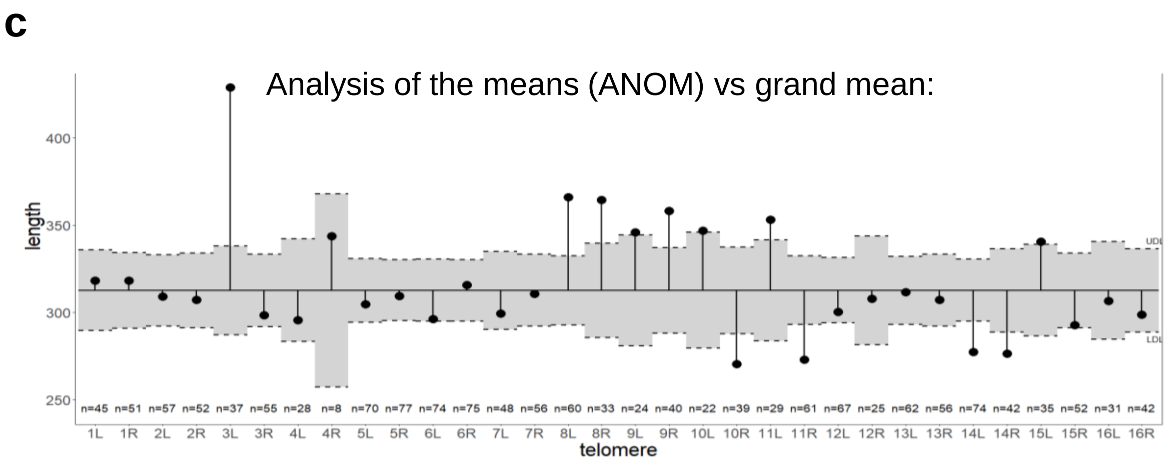

### Supplemental Figure 2

**a**

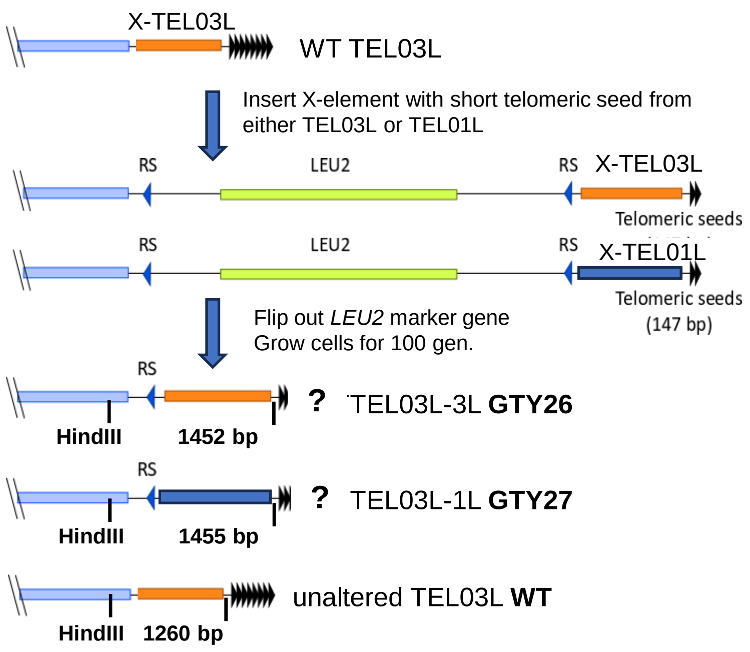


**b**

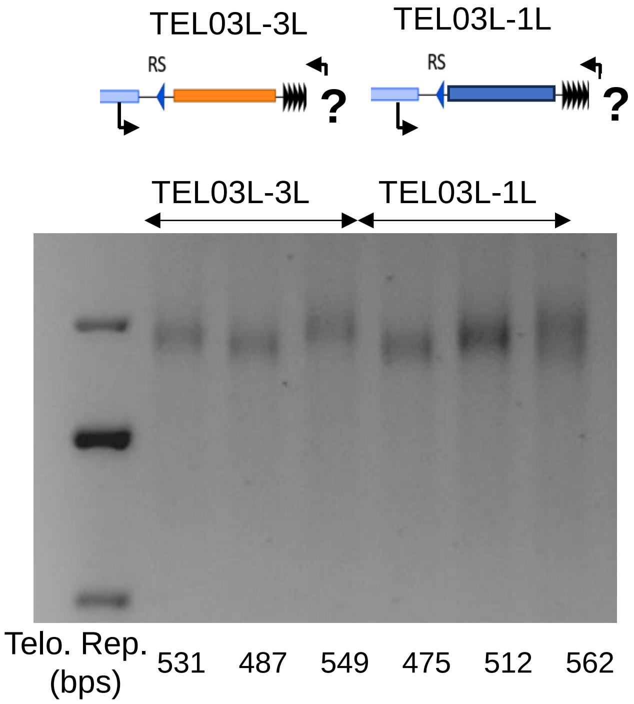

### Supplemental Figure 3

**a**

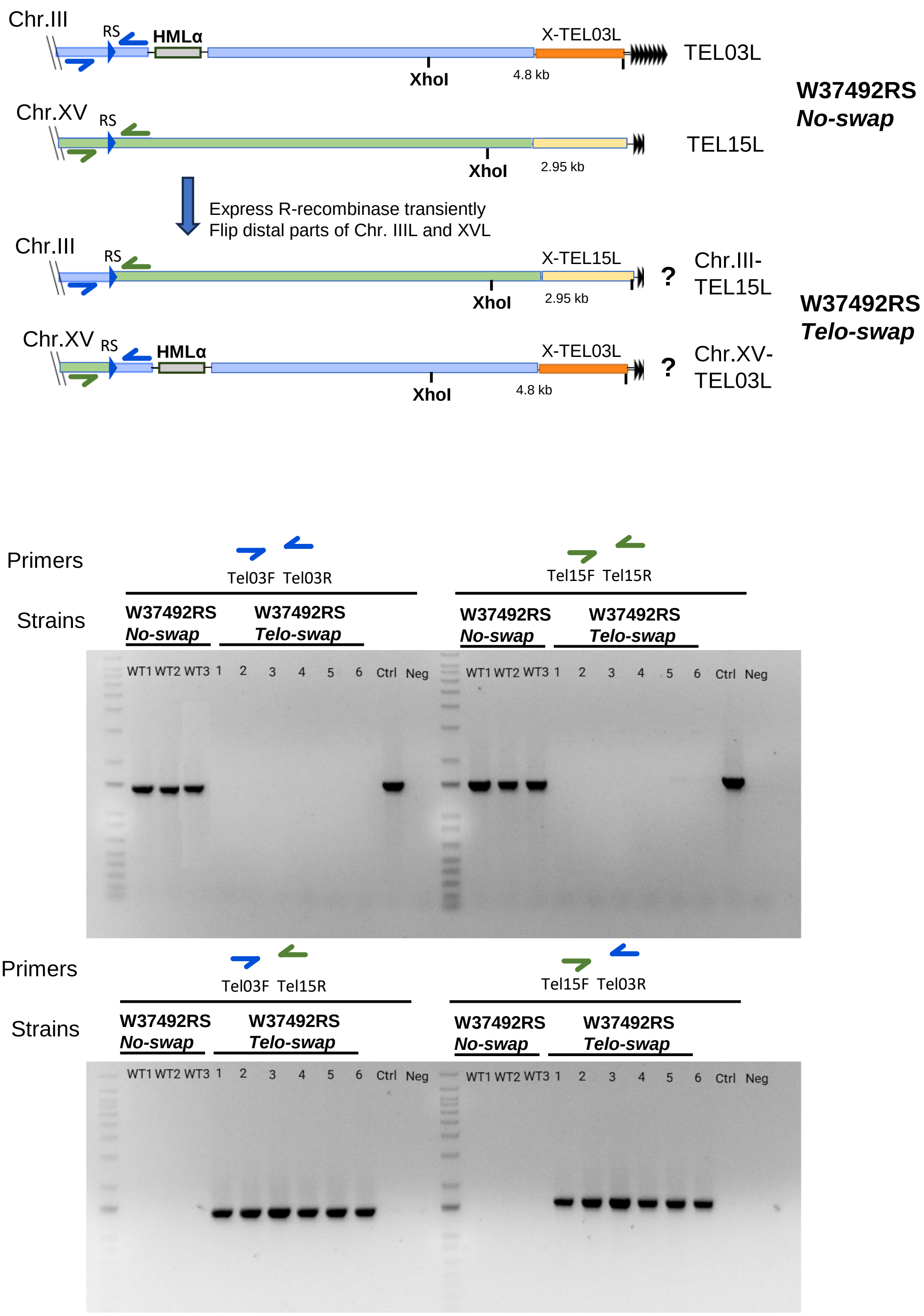


**b**

Supplemental Figure 3, continued

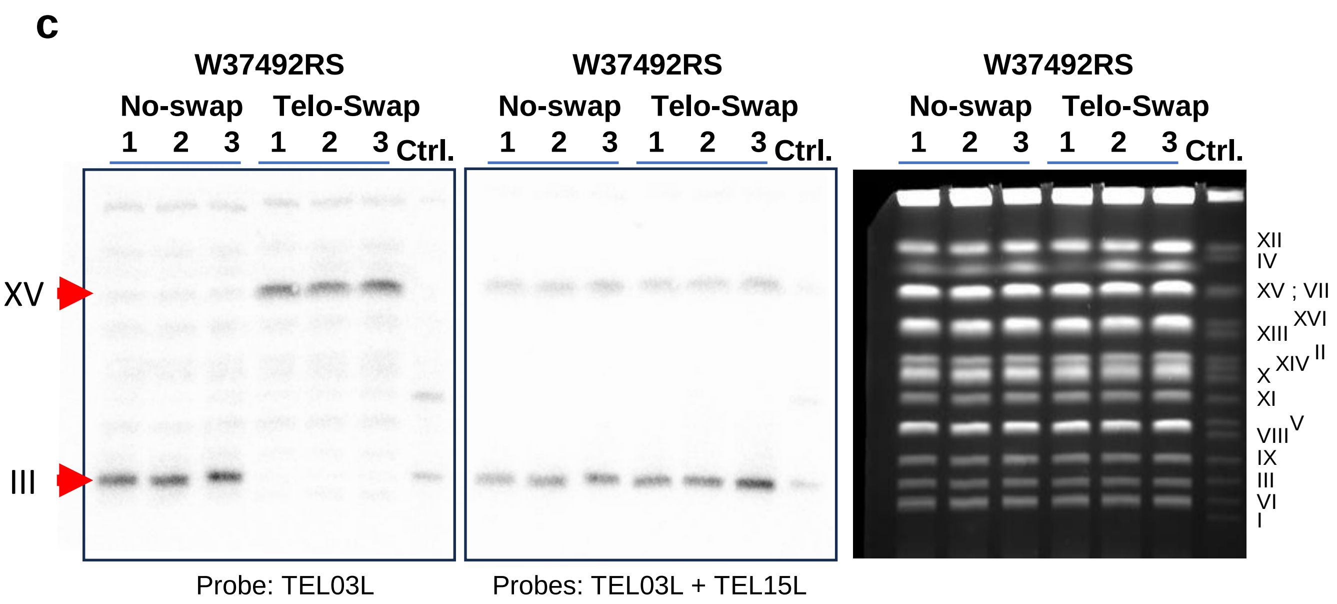

### Supplemental Figure 4

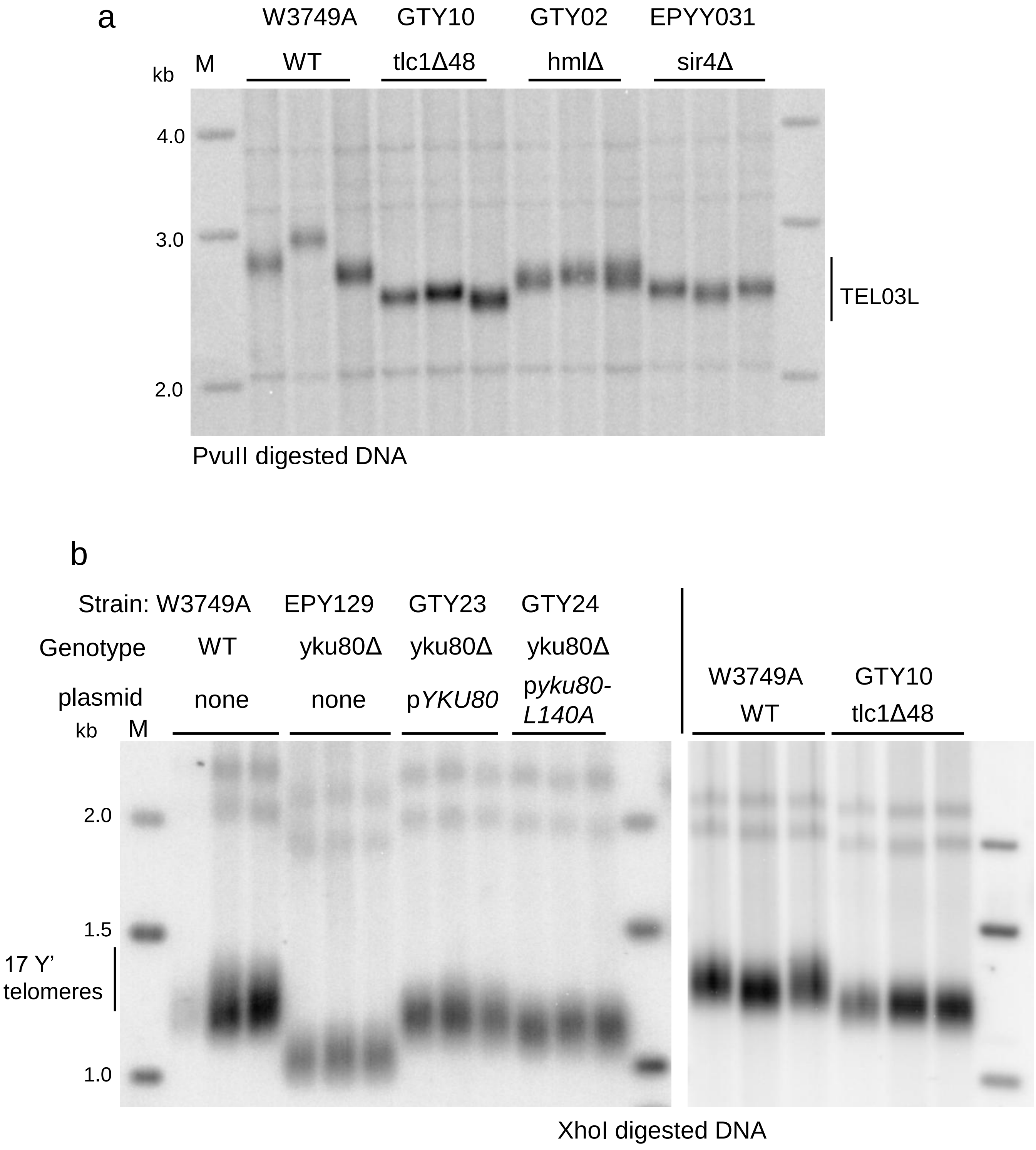

### Supplemental Figure 5

**a**

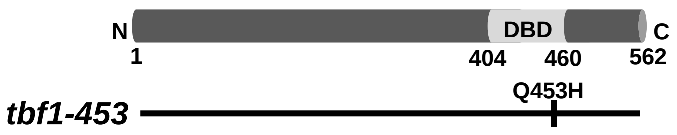


**b**

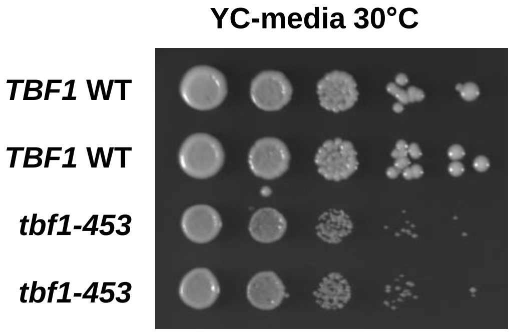


**c**

| WT vs *tbf1-453* 30°C; total RNA seq | |
|---|---|
| **RNAs**<br>**(+) = Tbf1 binding site in promoter**<br>**(-) = no Tbf1 binding site** | **Log2-ratio** |
| TBF1 (+) | 1.1 |
| PAF1 (+) | 0.9 |
| HAC1 (+) | 1.2 |
| PUF6 (-) | 0.6 |
| SNR190 (+) | -0.8 |
| SNR85 (-) | 1.7 |
| Telomeric mRNAs | |
| SIR1 | 1.3 |
| SIR2 | 0.4 |
| SIR3 | 1.5 |
| SIR4 | 1.1 |
| RIF1 | 1.2 |
| RIF2 | 0.4 |
| EST1 | 1.8 |
| EST2 | 1.7 |

**d**

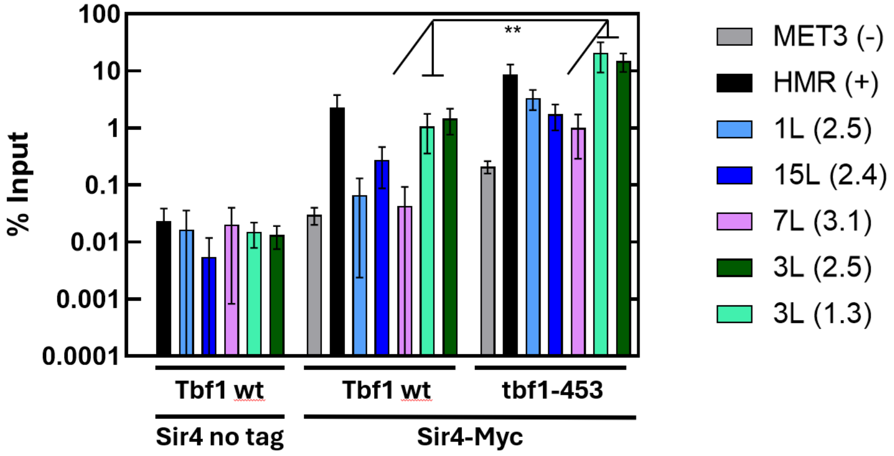

#### Supplemental Figure Legends:

##### Supplemental Figure 1:

***a***. TeloPCR analysis with DNA of 6 clones of WT W3749A. Telomeric DNA fragments amplified from TEL03L, TEL06R, and TEL11R with quantification of the tract lengths on the right (∗∗∗ for $p < 0.001$, t-test). ***b***. Telomere length distributions of each individual end in BY4741 WT cells (light blue plots) and BY4741 *sir3Δ* cells (light green plots) as obtained by Oxford Nanopore Sequencing. Lengths are reported in violin plots with a line for the mean length from WT. ***c***. Analysis of the means (ANOM) of each telomere against the grand mean of all telomere lengths in BY4741 WT cells was performed as reported [1].

##### Supplemental Figure 2:

***a.*** Schematic drawing of the construction of strains GTY26 and GTY27. GTY26 was constructed with a DNA fragment containing a *LEU2* gene flanked by RS sites, the TEL03L X-element and a short seed of telomeric repeat. GTY27 was constructed with a DNA fragment containing a *LEU2* gene flanked by RS sites, the TEL01L X-element and a short seed of telomeric repeat. After loss of the *LEU2* marker gene by recombination, cells were grown for 100 generations. The positions of the HindIII restriction sites on TEL03L in strains GTY26, GTY27 and the WT are indicated. ***b.*** TeloPCR of 3 clones of TEL03L-03L (GTY26) and 3 clones of TEL03L-1L (GTY27). Telomeric DNA fragments amplified from TEL03L and the repeat tract length of each is indicated below each lane.

##### Supplemental Figure 3:

***a.*** Schematic drawing of the construction of strains W37492RS *No-swap* and W37492RS *Telo-swap*. Starting strain W3749 was modified by inserting an RS recombination site 16.7 kb from the end of chromosome III and 13.5 kb from the end of chromosome XV using the same integration followed by flip out technology with a duplicated RS site as described in Suppl. Fig. 2a. The R-recombinase was then temporarily expressed from a plasmid and the status of the distal regions on chromosome IIIL and chromosome XVL verified by PCR with indicated by the primers. XhoI indicates location of that restriction enzyme site used for analysis in Fig. 3c. ***b.*** Verification PCR results for the Telo-swap on chromosome IIIL and chromosome XVL. WT1, WT2, and WT3 are 3 independent clones of the W37492RS *No-swap* strain, numbers 1-6 are 6 independent clones of the W37492RS *Telo-swap* strain and Ctrl. refers to the WT W3749 strain without integrated RS sites. Primer colors and direction are as in graph of Suppl. Fig. 3a. ***c.*** Pulsed field electrophoresis (CHEF gel) of whole chromosomes from 3 independent No-swap clones and 3 independent Telo-swap clones. Ctrl.: NEB control yeast chromosomes. Right: EtBr stained gel before blotting; Left: blot was hybridized to a TEL03L-specific probe; Middle: Blot was hybridized to both TEL03L- and Tel15L-specifric probes. Location of chromosomes III and XV on the blot is indicated on the left.

**Supplemental Figure 4:**

***a***. Southern blot of DNA from W3749A WT cells, cells with the *tlc1Δ48* allele (GTY10), cells with the *hmlΔ* allele (GTY02) and cells with the *sir4Δ* allele (EPY031). DNA was digested with PvuII and probed with a TEL03L unique probe. ***b***. Left: Southern blot with the same DNA as in Fig. 4a (right), but digested with XhoI and probed with a telomeric repeat fragment. Right: DNA of cells of the indicated genotypes as in Fig. 4a but digested with XhoI and probed with a telomeric repeat fragment.

**Supplemental Figure 5:**

***a.*** Detail of the single amino acid mutation in *tbf1-453* allele. ***b.*** Growth characteristics of cells with the *tbf1-453* allele vs WT. The four strains were derived from a microdissected heterozygous diploid strain. ***c.*** Selected results of mRNA seq analysis of total RNA derived from WT and *tbf1-453* cells. log2 ratios of the indicated genes are listed (See Data Availability section for access to all data). (+) indicates that the promoter of that gene contains a predicted Tbf1 binding site; (-) indicates that this gene does not contain such a site. ***d.*** Quantitative ChIP-PCR for Sir4 binding in telomere proximal areas of the indicated telomeres. Results from the procedure with untagged Sir4 or on the *MET3* locus served as negative controls (-); the *HMRa* locus (+) serves as a positive control. Primers are on the indicated telomeres at a distance from the X-element as indicated in brackets in kb. The differences between *TBF1* WT and *tbf1-453* on the indicated loci are significant with ** $p < 0.01$; t-test. For the other loci (not indicated) $p < 0.05$; t-test.

#### Supplemental Tables

##### Supplemental Table 1

A list of the yeast strains and their genotype used in this work.

| Strain | Genotype | References |
|---|---|---|
| W303 MATa | *MATa leu2-3,112 his3-11,15 trp1-1 ura3-1 ade2-1 can1-100* | [1] |
| W3749 MATa | *MATa can1-100 ura3-1 his3-11,15 leu2-3,112 trp1-1 bar1::LEU2* | [2] |
| W3749 MATα | *MATα can1-100 ura3-1 his3-11,15 leu2-3,112 trp1-1 bar1::LEU2* | [2] |
| BY4741 | *MATa his3Δ leu2Δ0 ura3Δ0* | [3] |
| BY4741 sir3Δ | *MATa his3Δ leu2Δ0 ura3Δ0 sir3Δ::kanMX* | This study |
| GTY11 (sir1Δ W3749A) | *MATa can1-100 ura3-1 his3-11,15 leu2-3,112 trp1-1 bar1::LEU2 sir1Δ::KMX* | This study |
| GTY12 (sir2Δ W3749A) | *MATa can1-100 ura3-1 his3-11,15 leu2-3,112 trp1-1 bar1::LEU2 sir2Δ::KMX* | This study |
| GTY13 (sir3Δ W3749A) | *MATa can1-100 ura3-1 his3-11,15 leu2-3,112 trp1-1 bar1::LEU2 sir3Δ::KMX* | This study |
| EPY031 (sir4Δ W3749A) | *MATa can1-100 ura3-1 his3-11,15 leu2-3,112 trp1-1 bar1::LEU2 sir4Δ::KMX* | [4] |
| JNY301 (rad52Δ W303A) | *MATa leu2-3,112 his3-11,15 trp1-1 ura3-1 ade2-1 can1-100 rad52Δ::LEU2* | This study |
| JNY302 (rad52Δ W303α) | *MATα leu2-3,112 his3-11,15 trp1-1 ura3-1 ade2-1 can1-100 rad52Δ::LEU2* | This study |

| | | |
|---|---|---|
| AHY001 (Est1-FRB W3749A) | *MATa ade2-1 trp1-1 leu2-3,112 his3-11,15 ura3 GAL psi+ tor1-1 fpr1D::NAT RPL13A-2xFKBP12::TRP1 Est1-FRB::KanMX* | This study |
| AHY005 (Est3-FRB W3749A) | *MATa ade2-1 trp1-1 leu2-3,112 his3-11,15 ura3 GAL psi+ tor1-1 fpr1D::NAT RPL13A-2xFKBP12::TRP1 Est3-FRB::KanMX* | This study |
| GTY26 (TEL3L(WT XCR)) | *MATa leu2-3,112 his3-11,15 trp1-1 ura3-1 ade2-1 can1-100 TEL03L(WT XCR)::LEU2* | This study |
| GTY27 (TEL3L (TEL01L XCR)) | *MATa leu2-3,112 his3-11,15 trp1-1 ura3-1 ade2-1 can1-100 TEL03L( Tel01L XCR)::LEU2* | This study |
| GTY02 (HmlΔ W3749A) | *MATa can1-100 ura3-1 his3-11,15 leu2-3,112 trp1-1 bar1::LEU2 hmldel::ura3* | This study |
| GTY19 (HmlΔ W3749A) | *MATa can1-100 ura3-1 his3-11,15 leu2-3,112 trp1-1 bar1::LEU2 hmldel::TRP* | This study |
| GTY20 (HmrΔ W3749A) | *MATa can1-100 ura3-1 his3-11,15 leu2-3,112 trp1-1 bar1::LEU2 hmrdel::TRP* | This study |
| GTY10 (tlc1Δ48 W3749A) | *MATa can1-100 ura3-1 his3-11,15 leu2-3,112 trp1-1 bar1::LEU2 tlc1Δ48* | This study |
| EPY129 (yku80Δ W3749A) | *MATa ade2-1 can1-100 ura3-1 his3-11,15 trp1-1 leu2-3,112 bar1Δ::LEU2 yku80Δ::URA3* | This study |
| GTY23 (EpY129 + pJP7c) | *MATa ade2-1 can1-100 ura3-1 his3-11,15 trp1-1 leu2-3,112 bar1Δ::LEU2 yku80Δ::URA3 + pJP7c* | This study |
| GTY24 (EpY129 + pJP7-L140A) | *MATα A ade2-1 can1-100 ura3-1 his3-11,15 trp1-1 leu2-3,112 bar1Δ::LEU2 yku80Δ::URA3 + pJP7c-L140A* | This study |
| ELY268-6b (Tbf1-453 W4749A) | *MATa can1-100 ura3-1 his3-11,15 leu2-3,112 trp1-1 bar1::LEU2 tbf1-453::NatMX* | This study |
| CLY05 (ELY268-6b Sir2Δ) | *MATa can1-100 ura3-1 his3-11,15 leu2-3,112 trp1-1 bar1::LEU2 tbf1-453::NatMX sir2Δ::KMX* | This study |

| | | |
|---|---|---|
| CLY06 (ELY268-6b Sir3Δ) | *MATa can1-100 ura3-1 his3-11,15 leu2-3,112 trp1-1 bar1::LEU2 tbf1-453::NatMX sir3Δ::KMX* | This study |
| GTY28 (ELY268-6b Sir4Δ) | *MATa can1-100 ura3-1 his3-11,15 leu2-3,112 trp1-1 bar1::LEU2 tbf1-453::NatMX sir4Δ::KMX* | This study |
| GTY31 (Rpd3Δ W3749A) | *MATa ade2-1 can1-100 ura3-1 his3-11,15 trp1-1 leu2-3,112 bar1Δ::LEU2 Rpd3Δ::trp1* | This study |
| ELY269 2-c (Sir4myc) | *W303/W3749a Sir4-13Myc::KanMX* | This study |
| ELY269 2-a (tbf1453 sir4myc) | *W303/W3749a tbf1-453::NatMX Sir4-13Myc::KanMX* | This study |
| EDY06 (RS-URA3-RS-TEL03L) | *MATα can1-100 ura3-1 his3-11,15 leu2-3,112 trp1-1 bar1::LEU2 RS-URA3-RS-TEL03L* | This study |
| EDY07 (RS-URA3-RS-TEL15L) | *MATa can1-100 ura3-1 his3-11,15 leu2-3,112 trp1-1 bar1::LEU2 RS-URA3-RS-TEL15L* | This study |
| EDY10 (RS-TEL15L-RS-TEL03L) | *MAT can1-100 ura3-1 his3-11,15 leu2-3,112 trp1-1 bar1::LEU2 TEL15L-Chr.III TEL03L-Chr.XV* | This study |

**Supplemental Table 2**

List of all primers and their respective use in this work.

| Name | Sequence (5′ to 3′) | Purpose |
|---|---|---|
| EST1_CTer Tag_F | TGATGAGGACATCACCGTCCAAGTGCCA GATACTCCTACT-cggatccccgggttaattaa | Cloning AHY001 |
| EST1_CTer Tag_R | TTTCATATTATGATTTTTTCCCTCACCATTAC TTGTTCTC-gaattcgagctcgtttaaac | Cloning AHY001 |
| EST3_CterT ag_F | CGGATCGTTAAGTACTTTCCCATTTGTATA TAAATATTTA-cggatccccgggttaattaa | Cloning AHY005 |
| Est3_CTerTa g_R | TAACTCTCTCACACTTATAAAATATCGAGC CTGCAGAAGG-gaattcgagctcgtttaaac | Cloning AHY005 |
| HML-F | GAGCTCATCTAGAGCCTTACGAAG | Cloning GTY19 |
| HML-R | ATACCTCTAACTTAGAATGTTTCAGC | Cloning GTY19 |
| HMR-F | AACATATAGAAGGGTCCAATAAACTTAC | Cloning GTY20 |
| HMR-R | CAGATGCGGAATTGGTGAATTTTTA | Cloning GTY20 |
| GT-TLC1-3 | tcatgcaggcctcagaaatt | GTY10 selection |
| GT-TLC1-4 | cgagaaaaaaataaacagcgaact | GTY10 selection |
| Q453H mut FOR | aaatttgaagaacaggacgcatgtacaactgaaagataa agcc | Cloning ELY268-6b |
| Q453H mut REV | ggctttatctttcagttgtacatgcgtcctgttcttcaaattt | Cloning ELY268-6b |
| TEL03L-P1F | AGCTTTCATCATTCGCGCTGA | Probing (DNA fragment specifc for TEL03L) and qPCR |
| TEL03L-P1R | CGTCAACAGGTTATGAGCCCT | Probing (DNA fragment specifc for TEL03L) and qPCR |
| tel15Lp1-F | TCGCGATGCCAACAAAATTC | Probing (DNA fragment specifc for TEL03L) and qPCR |

| | | |
|---|---|---|
| tel15Lp1-R | TCGGCGTTCCATATCGACA | Probing (DNA fragment specifc for TEL03L) and qPCR |
| G tails | GGGGGGGGGGGGGGGGGG | TeloPCR |
| Tel06R | TAAAGGAATCCCCAGAGACCTC | TeloPCR |
| Tel03L | AGCTTTCATCATTCGCGCTGA | TeloPCR |
| Tel11R | AATCCAATAAACTTACTACAATATGACATATAAG | TeloPCR |
| TEL03L-P2F | GGGTGGTTGTTTTACGTAGATC | qPCR |
| TEL03L-P2R | GAAGTACAAAGCTCGCCTTG | qPCR |
| Probe07L-F | TTAGGAAACACTACCCTATTCATATTCAAC | qPCR |
| Probe07L-R | GTAGAAAGGACATAAATAGTTGACGTG | qPCR |
| Tel1Lprobe-F | TAGACAATAAGCTTCTGTACGAGG | qPCR |
| Tel1Lprobe-F | TGGAGATGCTCTTGTTTCTGAA | qPCR |
| TEL03L Seq For | GGAGCCGCCACAAGCAATAATTATCACAATG | Telo swap verification |
| TEL15L Seq R | TGATGCTCTTATCAGAAGCTTAGAACTACAGAGAG | Telo swap verification |
| TEL15L Seq F | CGTCTGAATTGAACGCTCATTCAGAAGCTTATTG | Telo swap verification |
| TEL03L Seq R | TCCACAGATCTAATTGCAAGATAGCCTCTTGCG | Telo swap verification |

**Supplemental Table 3**

List of plasmids used with their primary features.

| Plasmid | Description | Reference |
|---|---|---|
| pLK2 | | Brewer/Raghuraman Labs; Uni. of Washington |
| pJH2039 | HMLΔ::NatR | [5] |
| T3TD | TEL03L subtelomeric region, XCR wt, X core, telomeric repeats with LEU2 flanked by RS site. | This study |
| pAH01 | TEL03L subtelomeric region, XCR Tel01L, X core, telomeric repeats with LEU2 flanked by RS site. | This study |
| pGT02 | HMLΔ::TRP1 | This study |
| pGT03 | HMRΔ::TRP1 | This study |
| pRS306-tlc1Δ48s | 48-nt deletion from nucleotides 288-355 of Tlc1. | [6] |
| pJP7C | YKU80-Mycx2-Hisx10, CEN, TRP1 | [7] |
| pJP7C-L140A | YKU80-L140A-Mycx2-Hisx10, CEN, TRP1 | [7] |
| pEP21A | Bacterial RS Recombinase expressed from a gal promoter | This study |
